# Short 5′ UTR enables optimal translation of plant virus tricistronic RNA via leaky scanning

**DOI:** 10.1101/2021.05.14.444105

**Authors:** Yuji Fujimoto, Takuya Keima, Masayoshi Hashimoto, Yuka Hagiwara-Komoda, Naoi Hosoe, Shuko Nishida, Takamichi Nijo, Kenro Oshima, Jeanmarie Verchot, Shigetou Namba, Yasuyuki Yamaji

## Abstract

Regardless of the general model of translation in eukaryotic cells, a number of studies suggested that many of mRNAs encode multiple proteins. Leaky scanning, which supplies ribosomes to downstream open reading frames (ORFs) by read-through of upstream ORFs, is the most major regulatory mechanism to translate polycistronic mRNAs. However, the general regulatory factors controlling leaky scanning and their biological relevance have rarely been elucidated, with exceptions such as the Kozak sequence. Here, we have analyzed the strategy of a plant RNA virus to translate three movement proteins from a single RNA molecule through leaky scanning. The *in planta* and *in vitro* results indicate that significantly shorter 5′ UTR of the most upstream ORF promotes leaky scanning, potentially finetuning the translation efficiency of the three proteins in a single RNA molecule to optimize viral propagation. Moreover, in plant endogenous mRNAs, we found that shorter UTRs were more frequently observed in uORFs of polycistronic mRNAs. We propose that the promotion of leaky scanning induced by a <u>sh</u>ort 5′ UTR (LISH), together with the Kozak sequence, is a conserved gene regulation mechanism not only in viruses but also in eukaryotes.

## Introduction

According to the general model of translation in eukaryotic cells, the translation machinery recognizes a single open reading frame (ORF) in a monocistronic mRNA. This recognition process has three steps: initiation, elongation, and termination. The first step, translation initiation, is a highly regulated process dependent on the 5′ cap structure of mRNA (1). The 5′ cap structure is recognized by the cap-binding protein eukaryotic translation initiation factor 4E (eIF4E) followed by recruitment of the 43S preinitiation complex (PIC) to the mRNA. Then the PIC scans the mRNA downstream from the 5′ end until the first appropriate initiation codon is recognized (2). Recognition of the initiation codon induces recruitment of the 60S ribosomal subunit to form the 80S ribosome complex, which transitions polypeptide synthesis referred to as the elongation step. Finally in the termination step, the ribosomes and synthesized proteins are released from the mRNA after stopping at the termination codon.

Increasing experimental evidence has demonstrated that many eukaryotic mRNAs have polycistronic structure: a single RNA molecule encoding multiple proteins. In fact, 44% of human mRNAs contain upstream ORFs (uORFs) located upstream of the start codons of annotated ORFs (3). Moreover, the start codon of uORF is more conserved among various mammalian species than would be estimated for neutral evolution, suggesting that this uORF plays a particular biological role (4, 5). Similar to animals, approximately 35% of mRNAs in the model plant *Arabidopsis thaliana* contain uORF, and half of them have multiple uORFs (6). Translation from polycistronic mRNA is accomplished mainly by read-through of an initiation codon referred to as leaky scanning. In leaky scanning, the PIC does not recognize the upstream AUG (uAUG) and reads past it, arriving at a downstream AUG (dAUG) to start the translation of the downstream ORF (dORF). Because the dAUG competes with the uAUG for the entry of PIC, increase of translation from the uAUG decreases translation from the dAUG, whereas decrease of translation from the uAUG increases translation from the dAUG.

The efficiency of AUG codon recognition by PIC depends on the nucleotide sequence context, and in particular, the optimal context is called the Kozak sequence (7). The context of the uAUG is an important regulatory factor in dORF expression because high efficiency of uAUG recognition promoted by a strong translation initiation sequence context disturbs the translation initiation from the dAUG. In fact, mutations in the translation initiation context of a uORF can result in serious physiological disorders at the organismal level, such as tumorigenesis in humans caused by natural variants in the Kozak sequence of a single gene (8). However, the regulatory mechanism of leaky scanning in eukaryotic cells is not fully understood.

Viruses are intracellular parasites that rely heavily on the host machinery for their propagation. To synthesize viral proteins, viruses retain a sophisticated mechanism to exploit the host translation system (9, 10). Presumably due to their limited genome size, RNA virus genomes are frequently polycistronic and they are expressed by multiple strategies. The genus *Potexvirus* includes a group of monopartite positive-sense, single-stranded RNA viruses that infect a wide range of plant species. The potexviral genome with a 5′ cap and 3’ poly (A) tail possesses five ORFs encoding RNA-dependent RNA polymerase (RdRp); triple gene block protein (TGBp) 1, 2, and 3; and coat protein (CP) (11, 12). TGBp1/2/3, which function as movement proteins (MPs) required for the intracellular movement of viruses in plant cells, are translated from three partially overlapping ORFs. Both the gene structure and the functions of TGBps are conserved in multiple genera of plant viruses other than potexviruses. Translation of these three proteins is considered to require two subgenomic RNAs (13) but the detailed mechanism of this translation system is not fully understood.

Here, we investigated the translation mechanism of TGBps and found that the three proteins are all translated from a single RNA molecule, sgRNA1, via leaky scanning. Through *in vitro*, *in planta*, and *in silico* analyses of the mechanism that regulates the translation of TGBps from sgRNA1 and the translation of *Arabidopsis thaliana* mRNAs, we showed that, in addition to the Kozak sequence, the length of the uORF 5′ UTR regulates leaky scanning. Thus, we propose a model named leaky scanning induced by a <u>sh</u>ort 5′ UTR (LISH) as a highly conserved translation regulatory mechanism, not only in viruses but also in their eukaryotic hosts.

## Material and methods

### Plant materials and growth conditions

*Nicotiana benthamiana* plants were maintained in a growth chamber with a 16-h light, 25°C/8-h dark, 20°C cycle throughout the assays. Detailed conditions for *Arabidopsis* cell culture were described previously (14). Detailed conditions for *Nicotiana* cell culture were described previously (15).

### Antibodies

Anti-PlAMV-TGBp1, -TGBp2, -TGBp3, -CP, -RdRp, and anti-PVX-CP antibodies were prepared as described previously (16–20). Other antibodies were raised in rabbits using purified peptides as antigens (Eurofins Genomics, Tokyo, Japan). Detailed information on antigens is shown in Supplementary Table 1.

### Plasmid construction

The sequences of all the primers used in this study are shown in Supplementary Table 3.

i. **Infectious cDNA clone.** Infectious GFP-expressing PlAMV (pPlAMV-GFP; 21,22) and PVX (pPVX-GFP) isolates were obtained as described previously. Infectious cDNA clones of PlAMV (pPlAMV; 21,23), PVX-OS strain (pPVX; 20), LoLV (pLoLV; 24), hydrangea ringspot virus (HdRSV; pHdRSV; 24), and white clover mosaic virus (WClMV; pWClMV; 24) were constructed as described previously. Infectious clones of pepino mosaic virus (PepMV; pPepMV) and PVM (pPVM) were constructed as follows. Full-length cDNA sequences of PepMV (PV-0716; Leibniz-Institut) and PVM (MAFF number 307027) were amplified using the PepMV1F/GRR and PVM1F/GRR primer sets, respectively. DNA fragments harboring the 35S promoter and nopaline synthase (NOS) terminator from the pCAMBIA1301 vector backbone with 5′- and 3′-terminal sequences of PepMV and PVM were amplified from pPPVOu binary vectors (25) using the GRF/PepMV35SR and GRF/PVM35SR primer sets, respectively. Full-length viral cDNAs were assembled with the corresponding PCR product consisting of the 35S promoter, NOS terminator, and pCAMBIA1301 vector backbone using a GeneArt Seamless Cloning and Assembly Kit (Thermo Fisher Scientific, Waltham, MA, USA) to generate pPepMV and pPVM.
ii. **Δsg1 mutant virus.** To construct the sgRNA1-deleted PlAMV mutant (Δsg1), several silent nucleotide substitutions were introduced into the sgRNA1 promoter while leaving the RdRp amino-acid sequence unchanged, according to a previous report (26). Partial viral sequences containing nucleotide substitutions and a binary vector fragment were amplified from pPlAMV using the 35SF/PlAMVdsg1R, PlAMVdsg1F/NOSR, and NOSF/35SR primer sets. The three fragments were assembled using a GeneArt Seamless Cloning and Assembly Kit. PlAMV-GFP-Δsg1 was constructed using the same primer sets as for PlAMV-Δsg1, with a distinct template, pPlAMV-GFP. LoLV-Δsg1 and PVM-Δsg1 were constructed as follows. Partial viral sequences were amplified from pLoLV or pPVM using the primer sets 35SF/LoLVdsg1R and LoLVdsg1F/NOSR or 35SF/PVMdsg1R and PVMdsg1F/NOSR, respectively. Fragments derived from each virus were assembled with the vector backbone generated from the NOSF/35SR primer set using a GeneArt Seamless Cloning and Assembly Kit. To generate PVX-Δsg1, we amplified a DNA fragment containing the nucleotide substitutions from pPVX using the OSdsg1F/OSdsg1R primer set. The amplified fragments were self-ligated using a Ligation-Convenience Kit (Nippon Gene, Tokyo, Japan). PVX-GFP-Δsg1 was constructed in the same manner as for PVX-Δsg1, with pPVX-GFP used as the template.
iii. **35S-Fluc, 35S-Rluc, and amino-acid substitution Rluc mutants.** 35S-Fluc was constructed as follows. The coding region of the *FLUC* gene was amplified via PCR using the 35S-fluc-1F and 35S-fluc-1653R primer set. The plasmid fragment was obtained from 35S-sGFP (27) using the 35S-down1F and 35S-up1R primers. The PCR-amplified *FLUC* gene was inserted downstream of the 35S promoter in the plasmid fragments using a GeneArt Seamless Cloning and Assembly Kit. 35S-Rluc was constructed as described previously (28). An amino-acid substitution Rluc series was constructed via PCR with 35S-Rluc and M1A as templates using the primer sets shown in Supplementary Table 4.
iv. **35S-sg1.** The sgRNA1 region of the PlAMV genome (21) was amplified via PCR using the 35S-4224F and 35S-6102R primers. The plasmid fragment was obtained from 35S-sGFP (27) using the 35S-down1F and 35S-up96R primers. The PCR-amplified sgRNA1 sequence was inserted downstream of the 35S promoter of the plasmid fragments using a GeneArt Seamless Cloning and Assembly Kit.
v. **sg1-TGBp1_Rluc, sg1-TGBp2_Rluc, sg1-TGBp3_Rluc, and sg1-TGBp1_Rluc-M14L.** The Rluc coding region was amplified from 35S-Rluc via PCR using the Rluc-1F and Rluc-936R primers. Plasmid fragments for sg1-TGBp1_Rluc, sg1-TGBp2_Rluc, and sg1-TGBp3_Rluc were obtained from 35S-sg1 using the primer sets Rluc-TGB13UF/35S-TGB15U-RlucR, Rluc-TGB23UF/TGB25U-RlucR, and Rluc-TGB33UF/TGB35U-RlucR, respectively. The PCR-amplified *RLUC* gene was inserted into each plasmid fragment using a GeneArt Seamless Cloning and Assembly Kit. The Rluc-M14L coding region was amplified from the M14L mutant via PCR using the Rluc-1F and Rluc-936R primers and inserted into the plasmid fragment for sg1-TGBp1_Rluc to construct sg1-TGBp1_Rluc-M14L.
vi. **Leader-sequence variants KS, Δstp, KZ(-3A)U, KZ(-3A)C, KZ(-3A)G, 5U5nt, 5U3nt, 5U1nt, dp5U, 5Usub1, and 5Usub2.** Fragments were amplified via PCR from 35S-sg1, sg1-TGBp1_Rluc, sg1-TGBp1_Rluc-M14L, sg1-TGBp2_Rluc, and sg1-TGBp3_Rluc using the primer sets shown in Supplementary Table 4. Amplified fragments were self-ligated using a Ligation-Convenience Kit.
vii. **TGBp2-KZ variants.** Fragments were amplified via PCR from 35S-sg1, sg1-TGBp2_Rluc, and sg1-TGBp3_Rluc using the primer sets shown in Supplementary Table 4. Amplified fragments were self-ligated using a Ligation-Convenience Kit.
viii. **GFP-containing variants.** The GFP coding region was amplified from the 35S-sGFP plasmid via PCR using the 35S-flag-GFP1F and GFP-5U-TGB1R primers. The plasmid fragments were obtained from 35S-sg1, sg1-TGBp2_Rluc, and sg1-TGBp3_Rluc using the TGB1F/35S-up96R primer set and ligated with the GFP fragment, resulting in GFP-sg1, GFP-sg1-TGBp2_Rluc, and GFP-sg1-TGBp3_Rluc, respectively. GFP-sg1-TGBp1_Rluc was constructed in the same manner using the 35S-flag-GFP1F/GFP-5U-Rluc1R primer set for the 35S-sGFP plasmid and the Rluc1F/35S-up96R primer set for sg1-TGBp1_Rluc. A GeneArt Seamless Cloning and Assembly Kit was used for ligation.
ix. **midAUG**. A partial sgRNA1 sequence that includes midAUG mutations (midAUGinsert) was synthesized by Eurofins Genomics (Tokyo, Japan). The midAUGinsert was ligated with plasmid fragments obtained via PCR from 35S-sg1, sg1-TGBp2_Rluc, and sg1-TGBp3_Rluc using the midAUGF/35S-up96R primer set. The sequence data are shown in Supplementary Table 3. The GFP coding fragment was ligated with plasmid fragments using a GeneArt Seamless Cloning and Assembly Kit.
x. **Leader-sequence variants of PlAMV-sgRNA1 and PVX-sgRNA1 for agroinoculation (PlAMV-sgRNA1-5U1nt, -3nt, -5nt, -7nt, and -10nt; PVX-sgRNA1-1nt, -4nt, -7nt, and -10nt).** The sgRNA1 sequences of pPlAMV and pPVX were amplified via PCR using the primer sets shown in Supplementary Table 4. Plasmid fragments were obtained from pPlAMV using the GRF/35SR primer set. The fragments were ligated using a GeneArt Seamless Cloning and Assembly Kit.
xi. **Leader-sequence variants of PVX-GFP for agroinoculation (PVX-GFP-5U4nt and -10nt).** The sgRNA1 sequence was amplified from pPVX-GFP via PCR using the primer sets shown in Supplementary Table 4. The plasmid fragments were obtained from pPVX-GFP using the PVXTGB1F/35SR primer set. The fragments were ligated using a GeneArt Seamless Cloning and Assembly Kit.

### Inoculation with viral isolates

Agroinoculation of PlAMV-GFP, PVX-GFP, PlAMV, PVX, LoLV, PepMV, WClMV, HdRSV, and PVM was performed as described previously (29). The leaf area with a GFP signal was quantitated using ImageJ v1.40 software [National Institutes of Health (NIH), Bethesda, MD, USA].

### qRT-PCR

Total RNA was extracted from transfected protoplasts using Sepasol-RNA I solution (Nacalai Tesque, Kyoto, Japan). Total RNA was subjected to DNase I treatment (Roche, Basel, Switzerland) followed by reverse transcription using a High Capacity cDNA Reverse Transcription Kit (Thermo Fisher Scientific). For qRT-PCR, viral RNA was amplified using a Thermal Cycler Dice Real Time System (TaKaRa, Shiga, Japan) with SYBR Premix Ex Taq II (TaKaRa). We used specific primers, RlucF and RlucR for *Rluc* mRNA, and PVXrtF and PVXrtR for PVX RNA. The accumulation of PVX RNA was normalized to *Rluc* Mrna to assess the activity levels and transformation efficiency in the protoplasts.

### Northern blot analysis

Total RNA (1 µ g) was analyzed using the digoxigenin (DIG) system (Roche). A DIG-labeled probe for PlAMV RNA detection was produced as described previously (16). Probes for PVX, LoLV, and PVM RNA detection were transcribed with T7 RNA polymerase from the DNA fragments amplified from each viral cDNA clone using the primer sets PVX5435F/PVX-RT7, LoLV6611F/LoLVRT7, and PVM7531F/PVMRT7, respectively.

### Protoplast preparation and transfection

*Arabidopsis* suspension culture cells (30) were kindly provided by S. Hasezawa (University of Tokyo). Protoplast isolation from *Arabidopsis* suspension cells and transfection were performed as described by Abel and Theologis (31) with slight modifications. First, 20 mL of suspension cells was collected by centrifugation and incubated with 10 mL of enzyme solution [1% cellulase Onozuka RS, 0.1% pectolyase Y-23, 0.4 M mannitol, and 5 mM MES-KOH (pH 5.6) dissolved in cell culture] for 120 min at 25°C. The cells were filtered through 100-µm nylon mesh to separate the protoplasts and washed three times with W5 buffer [154 mM NaCl, 125 mM CaCl_2_, 5 mM KCl, 5 mM glucose, and 1.5 mM MES-KOH (pH 5.6)]. The separated protoplasts were incubated on ice for 30 min before transfection. The protoplasts were counted in a hemocytometer and prepared at a density of 5 × 10^6^ protoplasts/mL. The protoplasts were collected and resuspended in the same volume of MaMg solution [0.4 M mannitol, 15 mM MgCl_2_, and 5 mM MES-KOH (pH 5.6)]. Then, 300 µL of protoplast solution was mixed with plasmid DNA (10 µg for the luciferase assay or 30 µg for immunoblotting). Next, 300 µL of polyethylene glycol-CHS solution (0.4 M mannitol, 0.1 M CaCl_2_, and 40% polyethylene glycol #4000) was added to the protoplast-plasmid mixture, which was incubated for 30 min at room temperature. The transfected protoplasts were washed with 5 mL of W5 buffer, resuspended in 2 mL of W5 buffer, and incubated in the dark at 23°C until subsequent analysis.

Protoplast isolation from *Nicotiana* suspension cells and transfection were performed as following. First, 30 mL of suspension cells was collected by centrifugation and washed with 20mL 0.4M mannnitol solution. Cells were incubated with 10 mL of BY-2 enzyme solution [1% cellulase Onozuka RS, 0.1% pectolyase Y-23, 0.4 M mannitol, and adjusted to pH5.5 by adding HCl] for 120 min at 25°C. The cells were washed three times with W5 buffer. The separated protoplasts were incubated on ice for 30 min before transfection. The protoplasts were counted in a hemocytometer and prepared at a density of 2 × 10^6^ protoplasts/mL. The protoplasts were collected and resuspended in the same volume of MaMg solution. Then, 300 µL of protoplast solution was mixed with 30 µg plasmid DNA. Next, 300 µL of polyethylene glycol solution (0.4 M mannitol, 0.1 M Ca(NO_3_)_2_, and 40% polyethylene glycol #4000) was added to the protoplast-plasmid mixture, which was incubated for 30 min at room temperature. The transfected protoplasts were washed with 5 mL of W5 buffer, resuspended in 2 mL of W5 buffer, and incubated in the dark at 25°C for three days until RNA extraction.

### Immunoblotting

Agroinfiltrated leaves were harvested at 4 dpi, total protein was extracted using RIPA buffer [50 mmol/L Tris-HCl (pH 8.0), 150 mmol/L NaCl, 0.5 w/v% sodium deoxycholate, 0.1 w/v% sodium dodecyl sulfate (SDS), 1.0 w/v% NP-40, and 100 mM dithiothreitol (DTT)]. The transfected protoplasts were collected at 42 h post-infection (hpi), and total protein was extracted using buffer A [50 mM Tris-HCl (pH 7.5), 15 mM MgCl_2_, 120 mM KCl, 10 mM DTT, 1 tablet Complete Mini Protease Inhibitor Cocktail (Roche, Basel, Switzerland)/10 mL, and 20 w/w% glycerol]. The sample was centrifuged at 1000 × *g* and 4°C for 10 min, and the supernatant was centrifuged again at 2000 × *g* and 4°C for 10 min to remove cell debris. The collected supernatant was centrifuged at 30,000 × *g* and 4°C for 30 min to generate the supernatant fraction (S30) and the membrane-containing pellet fraction (P30). Protein samples were denatured in gel sample buffer [50 mM Tris-HCl (pH 6.8), 2% SDS w/v, 10% glycerol, 100 mM DTT] and separated on a 3–8% Tris-acetate NuPAGE gel (Thermo Fisher Scientific) for RdRp, a 4–12% Bis-Tris gel for TGB1 and CP, and a 12% Bis-Tris gel for TGBp2 and TGBp3 of each virus. After electrophoresis, proteins were blotted onto a polyvinylidene fluoride membrane and detected with Can Get Signal (TOYOBO, Osaka, Japan). The membrane was stained with Coomassie brilliant blue as a loading control.

### Luciferase assay

Transfected protoplasts were collected at 19 hpi, and luciferase protein was extracted using extraction buffer [0.1 M phosphate (pH 7.0) and 5 µM DTT]. Luciferase activity was measured using a dual-luciferase reporter assay system (TOYO B-Net, Tokyo, Japan) according to the manufacturer’s instructions.

### 5′ RACE

The TSS of PlAMV sgRNA was detected in 5′ RACE analysis using a GeneRacer Kit (Invitrogen, Carlsbad, CA, USA), following the manufacturer’s instructions. Pr5010R was used as a primer specific for sgRNA1 and sgRNA2, whereas Pr5603R was used as a primer specific for sgRNA3. The TSSs of PVX, HdRSV, PepMV, WClMV, LoLV, and PVM were identified via sequencing on a MiSeq sequencing platform (Illumina, San Diego, CA, USA) using a MiSeq Reagent Kit v2 (500 cycles).

### Analysis of *Arabidopsis* UTRs

Sequence information for mRNAs was obtained from the TAIR9 version of the *A. thaliana* Col-0 genome (http://www.arabidopsis.org/). We identified the first AUG triplets in each mRNA when scanning from the 5′ end and classified them into two groups in which the first AUG triplet was matched or unmatched to the start codon of the annotated protein. The perl script used for classification is shown in Supplementary Table 5. For each of the two classified groups, we analyzed the length of the sequence upstream of the AUG triplet located at the 5′ end.

## Results

### All three TGB proteins are translated mainly from sgRNA1

Although TGBp2 and TGBp3 of potexviruses have been suggested to be translated from sgRNA2 (13), we did not find any clear signal corresponding to deduced sgRNA2 in northern blot analysis of *Nicotiana benthamiana* or *Arabidopsis thaliana* plants infected with the potexvirus plantago asiatica mosaic virus (PlAMV) (16; Fig. 1a, WT lane). Previous attempts by other research groups did not detect any clear putative sgRNA2 signal for other potexviruses (32, 33). To determine the transcription start sites (TSSs) of sgRNAs presumably transcribed from PlAMV genomic RNA, we used the 5′-rapid amplification of cDNA ends (5′-RACE) method with two gene-specific primers. The TSSs of sgRNA1 and sgRNA3 were successfully mapped to two major sites, G4224 and G5339, respectively, where the 5′ ends of all the cloned transcripts aligned (S. Fig. 1a, 1b). The same sites can be predicted to be TSSs based on consensus promoter sequences in viral genomic RNA (26). By contrast, no potential TSS was predicted for sgRNA2. The 5′ ends of cloned transcripts starting between the initiation codons of TGBp1 and TGBp2, which likely corresponded to the TSSs of sgRNA2, were consistently variable (S. Fig. 1c). These results indicated that a major TSS could not be defined for sgRNA2.

**Figure 1.**
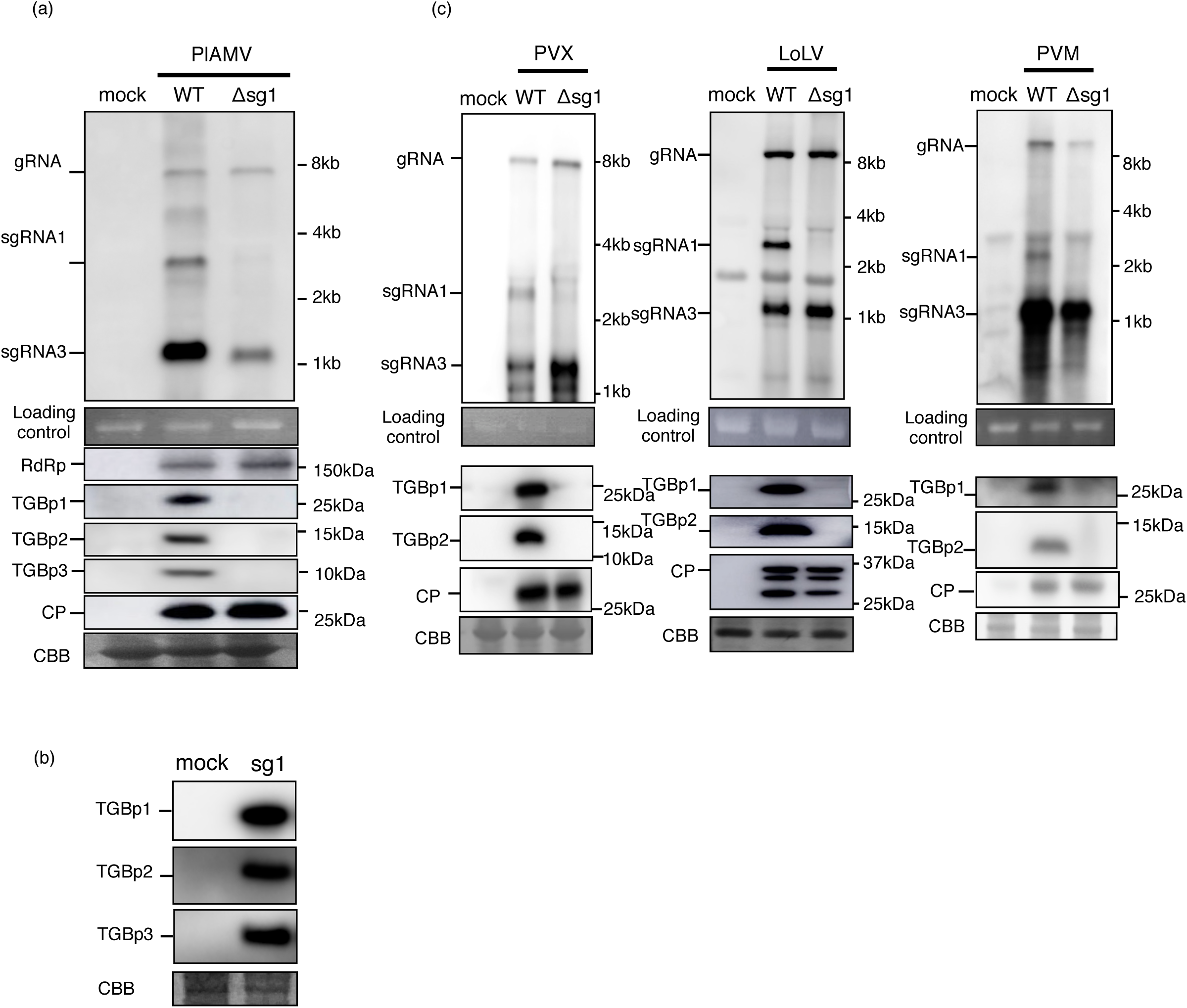
All flexivirus TGBps are translated from sgRNA1. (a) Northern blotting (upper window) and immunoblotting (lower windows) for the Δsg1 mutant of PlAMV. Total RNA and total protein samples extracted from non-inoculated plants were used as mock samples. (b) Immunoblotting of TGBps extracted from protoplasts transfected with PlAMV-sgRNA1 (sg1). A non-transfected sample was used as a mock sample. (c) Northern blotting (upper window) and immunoblotting (lower windows) for the Δsg1 mutant of PVX (genus *Potexvirus*, *Alphaflexiviridae*), LoLV (genus *Lolavirus*, *Alphaflexiviridae*), and PVM (genus *Carlavirus*, *Betaflexiviridae*). Total RNA and total protein samples extracted from inoculated plants were used as mock samples.

We hypothesized that the absence of sgRNA2 signals in northern blot analysis may be due in part to lower sgRNA2 accumulation. We reasoned that another viral RNA, most likely sgRNA1, participates in the translation of TGBp2/3. We prepared an sgRNA1-deleted PlAMV mutant (Δsg1), in which several nucleotide substitutions were introduced into the sgRNA1 promoter sequence, following the procedure in a previous report (26; S. Fig. 1d). To test whether the Δsg1 mutant produced TGBp1, TGBp2, and TGBp3, we inoculated the Δsg1 mutant into *N. benthamiana* leaves using the agroinoculation method. Total RNA extracted from the inoculated leaves at 1.5 days post-inoculation (dpi) was analyzed using northern blot hybridization. We confirmed that genomic RNA (gRNA) and sgRNA3 accumulated similarly as with wild-type (WT) PlAMV in Δsg1-inoculated leaves, but sgRNA1 did not (Fig. 1a). Immunoblot analysis revealed that TGBp2 and TGBp3, as well as TGBp1, were not detected in the Δsg1-inoculated leaves in which RdRp and CP accumulated to the same levels as in the WT (Fig. 1a). These results suggest that sgRNA1 is responsible for the translation of TGBp2 and 3 as well as TGBp1.

To further confirm the role of sgRNA1 in the production of TGBp2 and TGBp3 proteins, we examined whether TGBp2/3 were translated in sgRNA1-transfected protoplasts. Protoplasts isolated from *Arabidopsis* suspension culture cells were transfected with a plasmid expressing PlAMV sgRNA1 under the 35S promoter (sg1). Total protein extracted from the transfected protoplasts at 42 h was separated into soluble and insoluble fractions (S30 and P30, respectively). In sgRNA1-expressing protoplasts, TGBp1 was detected in both fractions, and both TGBp2/3 were specifically detected in the P30 fraction (Fig. 1b). This result together with the above finding indicates that sgRNA1 is a major template for the translation of TGBp2/3 in PlAMV.

To analyze whether the polycistronic nature of sgRNA1 is conserved in other groups of plant viruses with genomic structures similar to that of PlAMV, we examined the accumulation of TGBp1 and TGBp2 from sgRNA1 deletion mutants of viruses in the *Alphaflexiviridae* and *Betaflexiviridae* families. We introduced mutations into the sgRNA1 promoter sequences of potato virus X (PVX; genus *Potexvirus*) and lolium latent virus (LoLV; genus *Lolavirus*) in the family *Alphaflexiviridae*, and potato virus M (PVM; genus *Carlavirus*) in the family *Betaflexiviridae*. Total RNA and protein were extracted from leaves inoculated with each Δsg1 mutant. We confirmed that sgRNA1 was undetectable in all Δsg1 mutants, and there was no significant difference in the accumulation of gRNA, sgRNA3, or CP between the WT and Δsg1 mutants of all viruses tested, showing that these mutations impaired sgRNA1 expression but not viral multiplication (Fig. 1c). Consistent with the above findings for PlAMV, TGBp1 and TGBp2 were not detected via immunoblotting in the Δsg1 mutants of all viruses. These results indicate that the role of sgRNA1 in translating the TGBp1/2/3 proteins is conserved among several distinct virus species in *Alphaflexiviridae* and *Betaflexiviridae*.

### Translation of TGBp2/3 from sgRNA1 is not mediated by either an internal ribosome entry site (IRES) or reinitiation

Given the findings that TGBp1/2/3 are translated mainly from sgRNA1, we next analyzed the mechanism underlying the translation of TGBp1/2/3 from sgRNA1. Translation of dORFs in polycistronic mRNAs requires non-canonical mechanisms, such as leaky scanning (9). We hypothesized that the dORFs encoding TGBp2 and TGBp3 in sgRNA1 were regulated via leaky scanning for two reasons. First, according to the conventional potexviral TGBp expression model, leaky scanning regulates the translation of TGBp3 from sgRNA2 (13). Second, there is no AUG triplet between the TGBp1 and TGBp2 initiation codons in the PlAMV genome. This feature certainly favors leaky scanning to ensuring adequate translation of TGBp2/3 from sgRNA1.

To exclude the possibility that a non-canonical translation mechanism other than leaky scanning regulates the translation of TGBp2/3 from sgRNA1, we tested the involvement of two potential non-canonical translation mechanisms, IRES and re-initiation.

IRES is the structural RNA sequence that permits cap-independent translation initiation for the dORF by directly recruiting the translation apparatus including the PIC. We inserted a 40-nt Kozak-stem loop sequence (KS-sg1) to impair the migration of PIC scanning from the 5′ terminus (34,35; Fig. 2a). In addition, a green fluorescence protein (GFP)-coding sequence with a 21-nt 5′ leader sequence was fused to the 5′ end of sgRNA1 (GFP-sg1) to prevent majority of the PIC from the 5′ terminus from reaching to TGBp initiation codons by making the PIC recognize the AUG of GFP. If an IRES was located upstream of the start codon of TGBp2 to allow translation of TGBp2/3, the level of TGBp2/3 accumulation with KS-sg1 or GFP-sg1 would be similar to that with WT sgRNA1. Immunoblot analysis showed no accumulation of TGBp1/2/3 in *Arabidopsis* protoplasts transfected with KS-sg1 and GFP-sg1 (Fig. 2b), suggesting that sgRNA1 does not contain a functional IRES. The expression of GFP protein was confirmed via immunoblotting (Fig. 2b).

**Figure 2.**
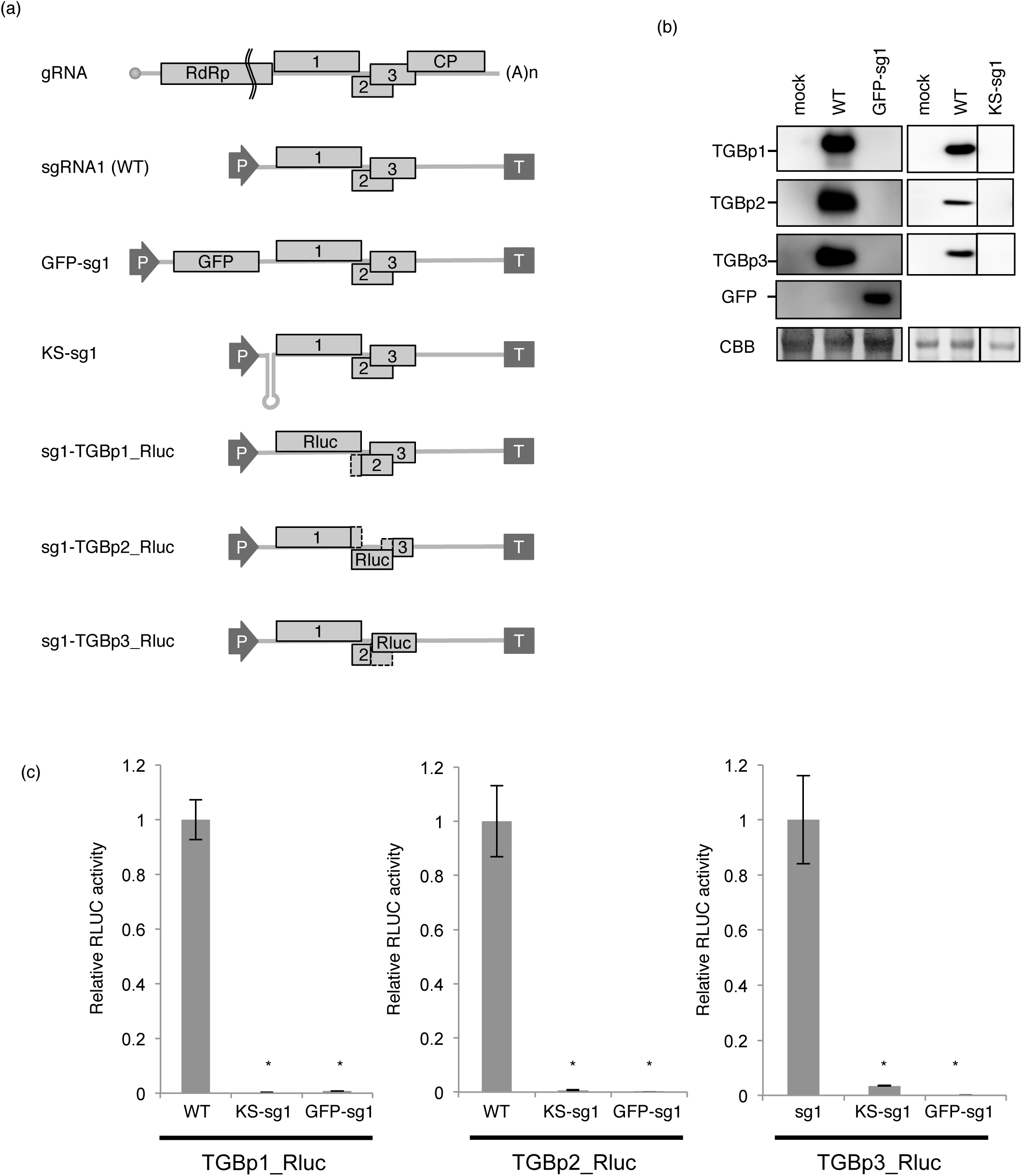
Translation of TGBp2/3 from sgRNA1 is not mediated by an IRES. (a) Schematic representation of PlAMV sgRNA1 cDNA constructs. Open box1, box2, and box3 represent TGBp1, TGBp2, and TGBp3 ORFs, respectively. The constructs contained full-length cDNA of sgRNA1 from a PlAMV-Pr isolate inserted between the cauliflower mosaic virus 35S promoter (P) and the NOS terminator (T). The 5′-terminal sequence of sgRNA1 is shown. A 40-nt Kozak stem loop sequence (19) was inserted between the sgRNA1 leader sequence and the TGBp1 initiation codon (KS-sg1). A *GFP*-coding sequence with a 21-nt 5′ leader sequence (AGCUGUACAAGUAAGAUUAAC) was fused to the 5′ end of sgRNA1 (GFP-sg1) to promote the recognition of the upstream *GFP* ORF by the majority of PIC from the 5′ terminus, so as to impair the migration of scanning PIC. (b) Immunoblot analysis of TGBp accumulation in protoplasts transfected with KS-sg1 and GFP-sg1 constructs. (c) Translational efficiency of TGBps in KS-sg1 and GFP-sg1 variants as analyzed using a dual luciferase assay. Rluc luminescence was normalized to Fluc luminescence. The mean values ± SDs from three independent experiments are shown. Asterisk indicates a significant difference compared with WT (Student’s t-test, P < 0.05).

To further evaluate the translational efficiency of TGBp1/2/3 in KS-sg1 and GFP-sg1 mutants, we performed a dual luciferase assay. We constructed modified KS-sg1, GFP-sg1, and WT sgRNA1 in which the coding region of TGBp1, 2, or 3 was replaced with the *Renilla* luciferase (*Rluc*) gene (Fig. 2a). The *Rluc*-containing sgRNA1 constructs were transfected into *Arabidopsis* protoplasts, and Rluc activity was normalized to firefly luciferase (Fluc) activity to assess the activity levels and transformation efficiency in the protoplasts. For all TGBp proteins, the relative Rluc activities of each mutant (KS and GFP constructs) were significantly lower than that of each WT construct (Fig. 2c), supporting the notion that sgRNA1 does not contain a functional IRES.

Next, to examine the possibility of the involvement of reinitiation, we replaced the UGA stop codon of TGBp1 with a GGA triplet (S. Fig. 2a). The frequency of reinitiation depends on the distance between stop codon of uORF and dAUG (36, 37). If TGBp2 and TGBp3 were translated via reinitiation, replacement of the TGBp1 stop codon would result in the clear separation between the initiation codon of TGBp2 and the end of the TGBp1 sequence, thereby reducing the level of TGBp2/3 expression. As expected, immunoblotting using anti-TGB1 antibody showed a band with a slightly higher molecular weight than that of the WT sg1-transfected protoplasts in the Δstp-transfected protoplasts, and a signal with a slightly lower molecular weight, presumably a degradation product, was also obtained (S. Fig. 2b). Both TGBp2 and TGBp3 accumulated in protoplasts transfected with Δstp to the same level as with WT-transfected samples (S. Fig. 2b). These results indicate that TGBp2 and TGBp3 are not translated from sgRNA1 via reinitiation.

### TGBp2/3 are translated via leaky scanning through the TGBp1 initiation codon

We tested our hypothesis that TGBp2 and TGBp3 are translated from sgRNA1 via leaky scanning of the TGBp1 initiation codon. In leaky scanning, the uAUG competes with the dAUG for translation initiation by ribosomes: the nucleotide context of the uAUG affects the efficiency of translation initiation and subsequently the continuation of PIC scanning to reach the dAUG. In dicots, the optimal sequence context for translation initiation is “RNN AUG G,” called the Kozak sequence, which includes a purine residue [R; guanine (G) or adenine (A)] at the −3 position, and a “G” at the +4 position relative to the first A in the initiation codon (38; Fig. 3a). The initiation codon of WT TGBp1 has an optimal Kozak sequence with A at the −3 position. The plasmids containing KZ(-3A) variants were transfected to *Arabidopsis* protoplasts and the accumulation of three TGB proteins were analyzed. When the A at −3 was replaced with the purine residue G [KZ(−3A)G], which produced another optimal sequence context, the accumulation of TGBp1, as well as TGBp2/3, was similar to WT (Fig. 3b). By contrast, when we introduced pyrimidine (U and C) to create a poor sequence context at the −3 position of the TGBp1 AUG codon, TGBp1 accumulation was reduced as expected (Fig 3b). Strikingly, TGBp2/3 accumulation increased in accordance with the decrease in TGBp1 accumulation (Fig. 3b). These immunoblot results can be explained if TGBp2/3 are translated via leaky scanning of the TGBp1 initiation codon in sgRNA1.

**Figure 3.**
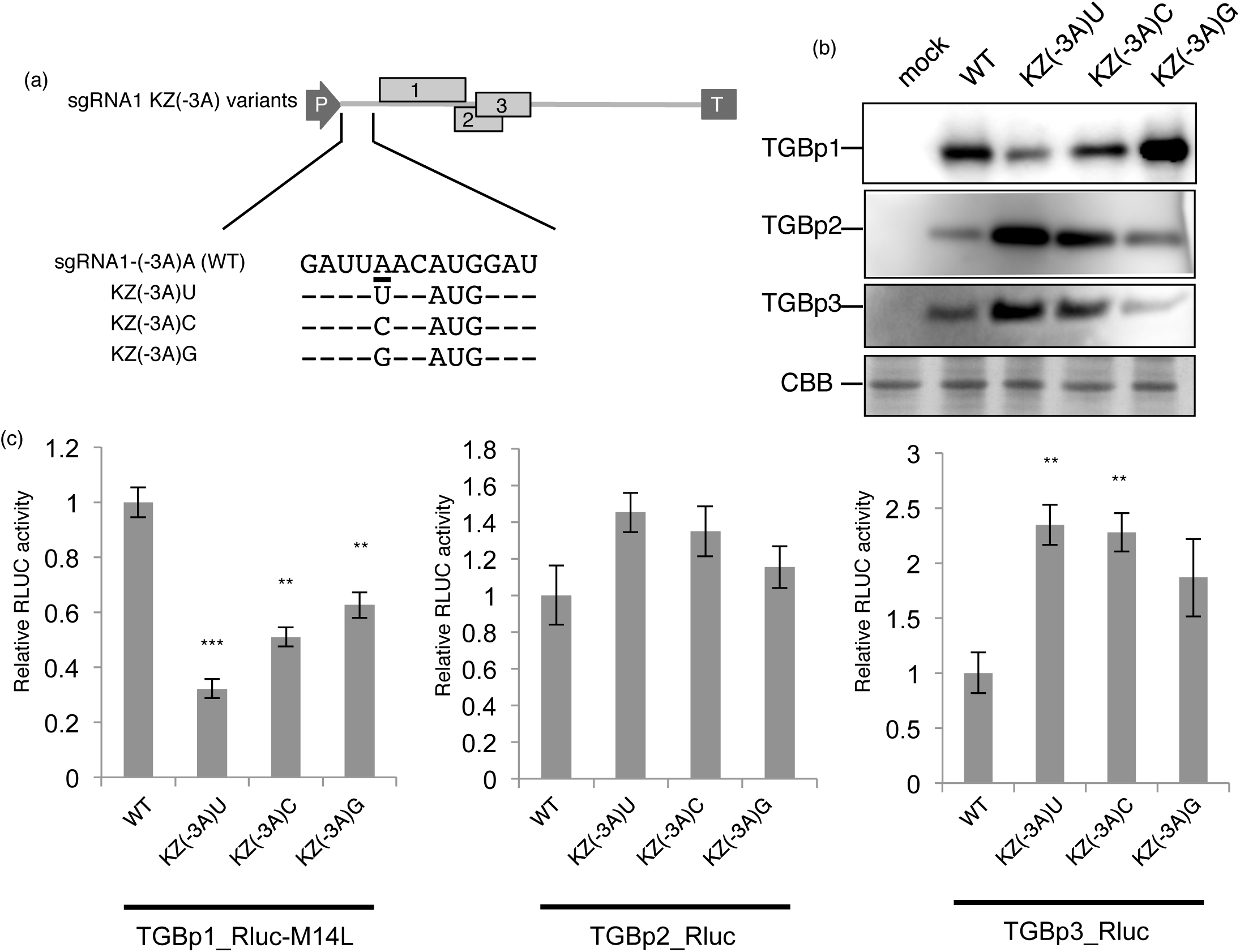
Translation of TGBp2/3 from sgRNA1 is mediated by leaky scanning. (a) KZ(−3A) variant constructs. The 5′-terminal sequence of sgRNA1 is shown. AUG is the initiation codon of TGBp1, and A at the −3 position (underlined) was changed to U, C, or G. Hyphens indicate that the nucleotide at that position was not substituted. (b) Immunoblot analysis of TGBp accumulation in KZ(−3A) variants . (c) Dual luciferase assay analysis of the translational efficiency of TGB proteins in KZ(−3A) variants. Rluc luminescence was normalized to Fluc luminescence. Mean values ± SDs from three independent experiments are shown. Asterisk indicates a significant difference compared with WT (Student’s t-test, double asterisk; P < 0.01, triple asterisk; P < 0.001).

Next, we used a dual luciferase assay to analyze how mutations in the TGBp1 Kozak sequence influence TGBp2/3 translation efficiency from sgRNA1 via leaky scanning. We replaced each TGBp-coding region with the *Rluc* gene in the above sgRNA1 Kozak −3 context variants and in WT sgRNA1. Note that we used an *Rluc* gene variant, Rluc-M14L, in which an in-frame second AUG triplet in the *Rluc* gene located 39 nt downstream of the canonical initiation codon was replaced with CUU to eliminate any possible interference from the potential translation of a partial gene product from the second AUG (S. Fig. 3). As shown in Figure 3C, the mutant with the purine substitution [sg1_KZ(−3A)G_TGBp1-Rluc-M14L], which had a potentially optimal Kozak sequence, exhibited higher translation efficiency from the TGBp1 initiation codon than did the mutants with the pyrimidine substitution [sg1_KZ(−3A)U_TGBp1-Rluc-M14L and sg1_KZ(−3A)C_TGBp1-Rluc-M14L]. Moreover, the mutant with the purine substitution exhibited lower TGBp2/3 translation efficiency than did the mutants with pyrimidine substitution. These data support the notion that the translational efficiency of TGBp2/3 in sgRNA1 is dependent on the nucleotide context of the TGBp1 initiation codon, potentially through leaky scanning. In line with the conventional TGBp expression model (13), we also confirmed that TGBp3 translation is actually mediated by leaky scanning of the TGBp2 initiation codon in sgRNA1 (S. Fig. 2c, d, e).

To further test the leaky scanning hypothesis, we introduced six AUG codons throughout the TGBp1 ORF by substituting nucleotides without changing the TGBp1 amino-acid sequence. In addition, the nucleotides located at −3 of each introduced AUG were replaced with G to satisfy the optimal Kozak sequence, unless the TGBp1 amino-acid sequence would be altered (S. Fig. 4a, midAUG). If TGBp2 and TGBp3 were translated from sgRNA1 by leaky scanning, insertion of several AUG codons preceding the AUG codons of TGBp2 and TGBp3 would reduce their translation efficiency by trapping the majority of PIC. The midAUG mutant and its *Rluc* derivatives were transfected into protoplasts, and expression of TGBp1/2/3 was analyzed via immunoblotting and luciferase assay. As expected, although immunoblotting showed that the accumulation of mutated TGBp1 did not differ from that of the WT, the levels of TGBp2 and TGBp3 and their translation efficiency as quantified by the luciferase assay were significantly reduced (S. Fig. 4b, c). These results also agree with our hypothesis that leaky scanning of the TGBp1 initiation codon regulates the translation of TGBp2/3 from sgRNA1.

As shown in Supplementary Figure 2c, d, and e, the translation of TGBp3 requires leaky scanning through the TGBp2 initiation codon. In summary, our results suggest that TGBp2 translation requires leaky scanning of the TGBp1 initiation codon, and TGBp3 translation requires leaky scanning of multiple upstream initiation codons in sgRNA1.

### The TGBp1 ORF 5′-leader sequence length regulates the efficiency of leaky scanning in sgRNA1

Although our results indicate that the nucleotide context of the TGBp1 initiation codon regulates the efficiency of leaky scanning, the nucleotide context of the TGBp1 initiation codon is inherently optimal. This made us postulate that an unknown factor promotes leaky scanning of the TGBp1 initiation codon, because TGBp2 and TGBp3 are translated sufficiently. As shown in Supplementary Figure 1, the TGBp1 leader in sgRNA1 is remarkably short (7 nt). Due to the conserved genomic structure among flexiviruses, we predicted that the mechanism of TGBp2/3 translation from sgRNA1 would be conserved among other flexiviruses. To examine whether this short leader sequence of sgRNA1 is widely conserved among potexviruses and related viruses, we determined the TSSs of the sgRNA1s of several potexviruses, a lolavirus, and a carlavirus. We found that the sgRNA1 5′-leaders of these viruses were all no more than 8 nt in length, supporting the notion that this short 5′-leader length in the translation of TGBp2/3 from sgRNA1 is conserved among these viruses (Fig. 4). Thus, we assumed that the short leader sequence of TGBp1 is another factor that regulates leaky scanning of sgRNA1.

**Figure 4.**
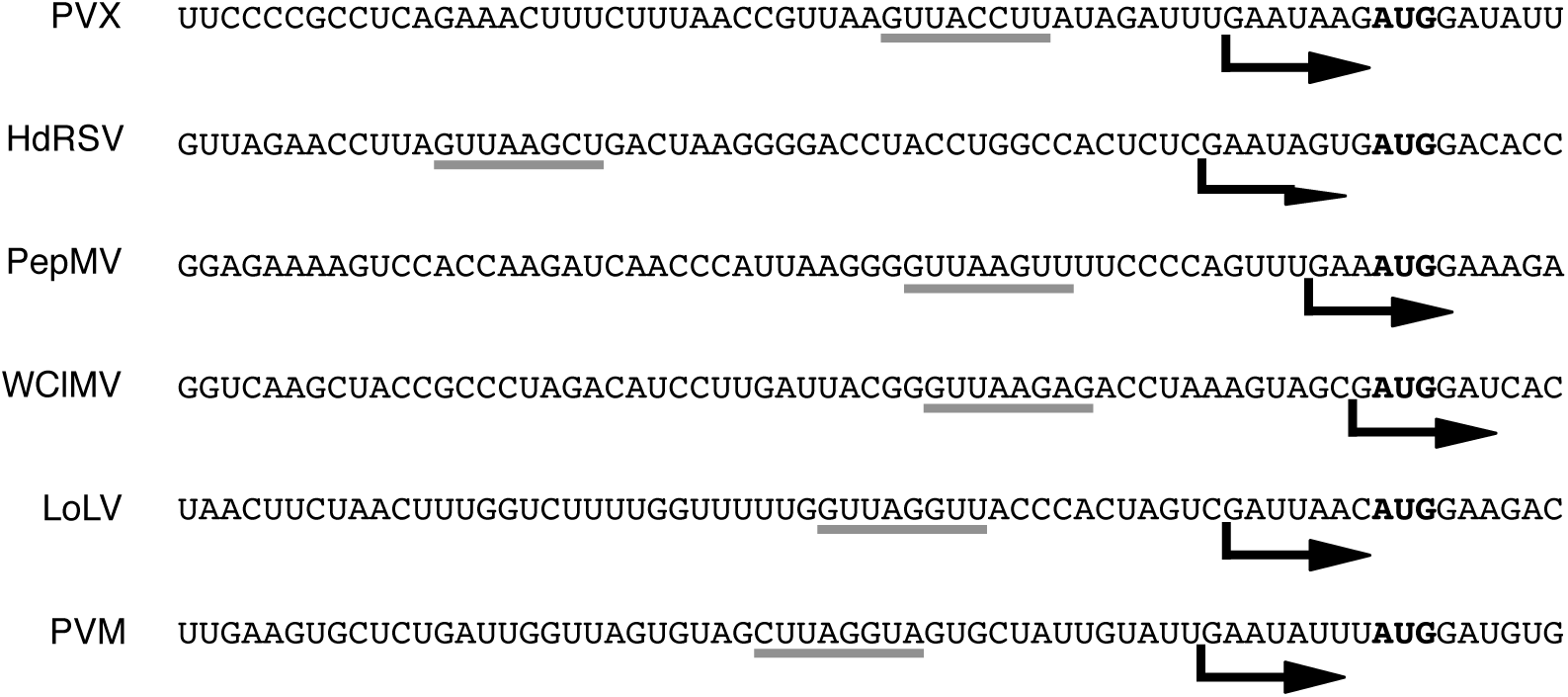
The short TGBp1 5′ leader sequence is highly conserved among several flexiviruses. PVX, HdRSV, WClMV, PepMV (genus *Potexvirus*, *Alphaflexiviridae*), LoLV (genus *Lolavirus*, *Alphaflexiviridae*), and PVM (genus *Carlavirus*, *Betaflexiviridae*) were analyzed. Deletion of sgRNA1 from PVX, LoLV (*Alphaflexiviridae*), and PVM (*Betaflexiviridae*) impairs TGB1/2 accumulation in a viral infection context, showing the relevance of sgRNA1 to the translation of all TGBps in other flexiviruses (Fig. 1). A survey of the TSSs of sgRNA1 of PVX, HdRSV, WClMV, PepMV, LoLV, and PVM showed that the length of the sgRNA1 5′ leader sequence was 7 nt in PVX and LoLV, 8 nt in HdRSV and PVM, 3 nt in PepMV, and 1 nt in WClMV. Putative promoter sequences are underlined. The arrow indicates the TSS of sgRNA1. Boldfaced AUG indicates the initiation codon of TGBp1. TSSs of each sgRNA1 are indicated by black arrows.

To test this hypothesis, we analyzed the levels of TGBp1/2/3 accumulation from PlAMV sgRNA1 variants whose leader lengths were shortened to 5, 3, or 1 nt (Fig. 5a). Immunoblotting revealed that TGBp1 levels gradually decreased as the leader length became shorter; by contrast, accumulation of TGBp2 and TGBp3 increased (Fig. 5b). Luciferase assay results also showed that the translational efficiency of TGBp2/3 increased in a stepwise manner as the 5′-leader sequence was shortened (7-nt length to 1-nt length), whereas that of TGBp1 decreased (Fig. 5c), indicating that shortening the leader sequence promoted the efficiency of leaky scanning. To further validate this hypothesis, we reasoned that extension of the sgRNA1 5′-leader sequence would reduce the efficiency of leaky scanning. To test this possibility, we constructed three sgRNA1 variants: dp5U with a tandemly duplicated WT leader sequence (14 nt), 5Usub1 with a heterologous long leader sequence (42 nt), and 5Usub2 with a heterologous GC-rich leader sequence with the same length as the WT (7 nt) (S. Fig. 5a). Immunoblotting and luciferase assay results showed that the translation levels of TGBp2/3 in protoplasts transfected with sgRNA1 variants with a long leader sequence (dp5U and 5Usub1) were lower than those produced from WT sgRNA1 (sg1), whereas the accumulation of TGBp1 did not differ (S. Fig. 5b, c). By contrast, 5Usub2 had the same leader length as the WT but a higher GC content, and was equivalent to the WT in terms of the accumulation levels of all TGBps (S. Fig. 5b, c). Taken together, these results indicate that leaky scanning induced by a <u>sh</u>ort 5′ UTR (LISH) enables ribosomes to efficiently translate TGBp2 and TGBp3.

**Figure 5.**
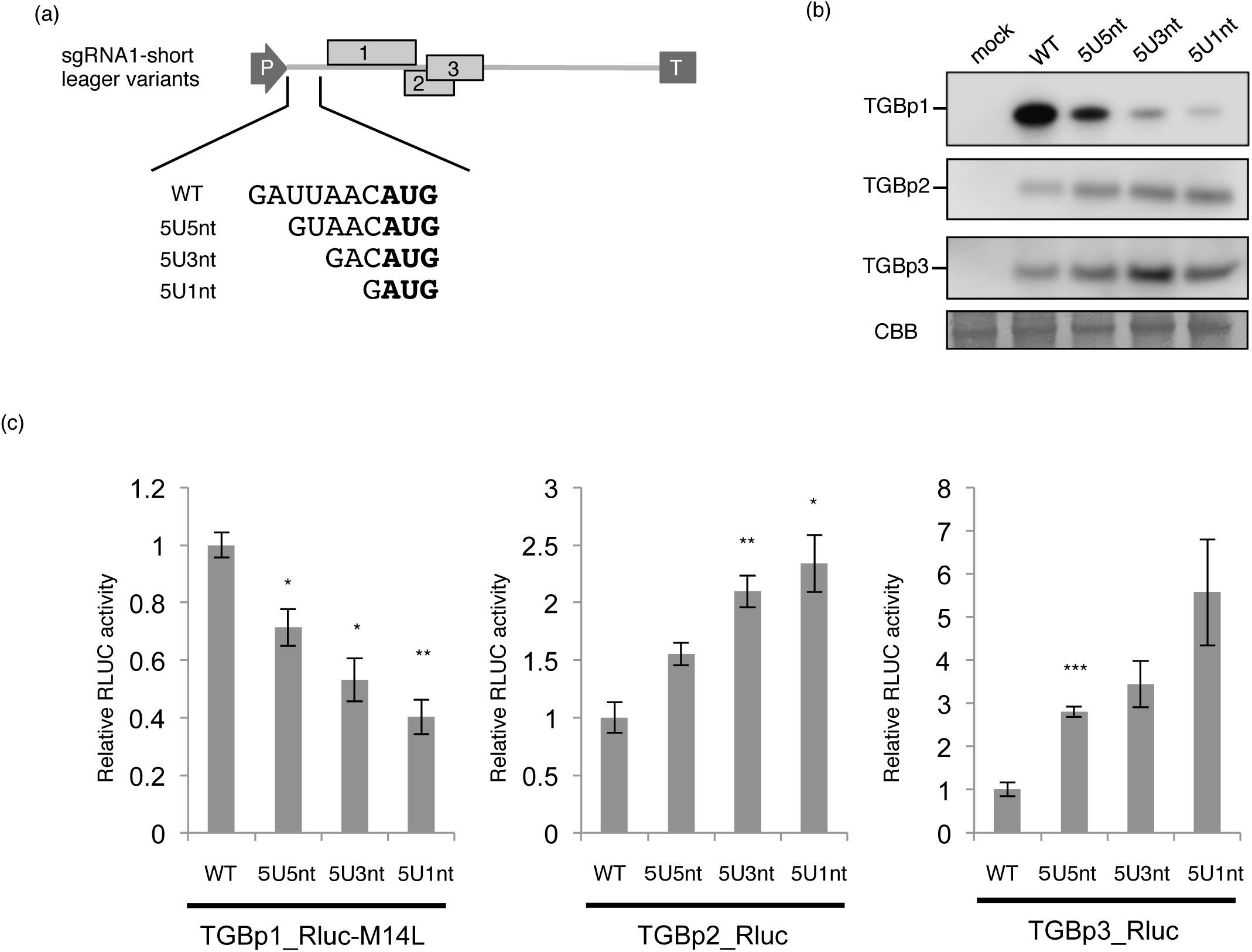
Short leader sequence increases the efficiency of leaky scanning. (a) Short leader variants constructs. The 5′-terminal sequence of sgRNA1 is shown. Boldfaced AUG indicates the initiation codon of TGBp1. (b) Immunoblot analysis of TGBp accumulation in short leader variants (c) Dual luciferase assay analysis of the translational efficiency of TGBp in short leader variants. Rluc luminescence was normalized to Fluc luminescence. Mean values ± SDs from three independent experiments are shown. Asterisks indicate a significant difference compared with WT (Student’s t-test, single asterisk; P < 0.05, double asterisk; P < 0.01, triple asterisk; P < 0.001).

### Leader sequence length is optimized for viral infection

Our results so far indicate that the amount of the three TGBps translation from sgRNA1 varies depending on the length of the leader sequence. Thus, to test whether this variation has biological relevance, we employed a *trans*-complementation system incorporating a GFP-labeled PVX mutant lacking sgRNA1 (PVX-GFP-Δsg1) in combination with transient transfection with PVX-sgRNA1 variants with various leader lengths (1, 4, 7, or 10 nt). Using GFP-tagged PVX, the efficiency of viral spread from infected cells to neighboring healthy cells (cell-to-cell movement) was quantified by measuring the area of GFP fluorescence (16). As shown in Supplementary Figure 6, GFP foci derived from the *trans*-complementation of sgRNA1 variants with different leader lengths were smaller than those resulting from WT leader-containing sgRNA1, suggesting that the alteration of LISH via the mutation of leader length modulated the cell-to-cell movement efficiency of PVX, likely due to an imbalance in the accumulation levels of TGBps (S. Fig. 6b, c). Similar results were obtained in PlAMV using a corresponding *trans*-complementation assay combining a GFP-labeled PlAMV mutant lacking sgRNA1 (PlAMV-GFP-Δsg1) and PlAMV-sgRNA1 variants with various leader lengths (1, 3, 5, 7, or 10 nt) (S. Fig. 7a). The efficiency of cell-to-cell movement of PlAMV-GFP increased as the leader sequence became longer, and, noticeably, the movement efficiency of the virus with a 10-nt leader was nearly the same as that of the WT (S. Fig. 7b, c). This may be because higher efficiency of cell-to-cell movement does not always maximize viral fitness (39).

Furthermore, we constructed PVX-GFP mutants to express sgRNA1 with various leader lengths in *cis* (Fig. 6a). The overlap of the RdRp coding region and the leader sequence of sgRNA1 prevented the construction of PlAMV variants without changing the amino-acid sequence of RdRp. We confirmed that the TSSs of sgRNA1 variants transcribed from these PVX mutants by RACE analysis were as intended. In addition, northern blotting and quantitative reverse transcription PCR (qRT-PCR) indicated that these mutations did not significantly affect viral RNA multiplication in protoplasts isolated from *Nicotiana* suspension culture cells (BY-2) (Fig. 6b, c). When GFP-tagged PVX variants expressing sgRNA1 with leader lengths of 4 or 10 nt were inoculated into *N. benthamiana*, the relative sizes of GFP fluorescence spots produced by the WT virus were significantly larger than those of mutants with shorter and longer leader sequences (Fig. 6d, e). Taken together, our results suggest that the native leader length optimizes the efficiency of LISH to achieve optimal cell-to-cell movement.

**Figure 6.**
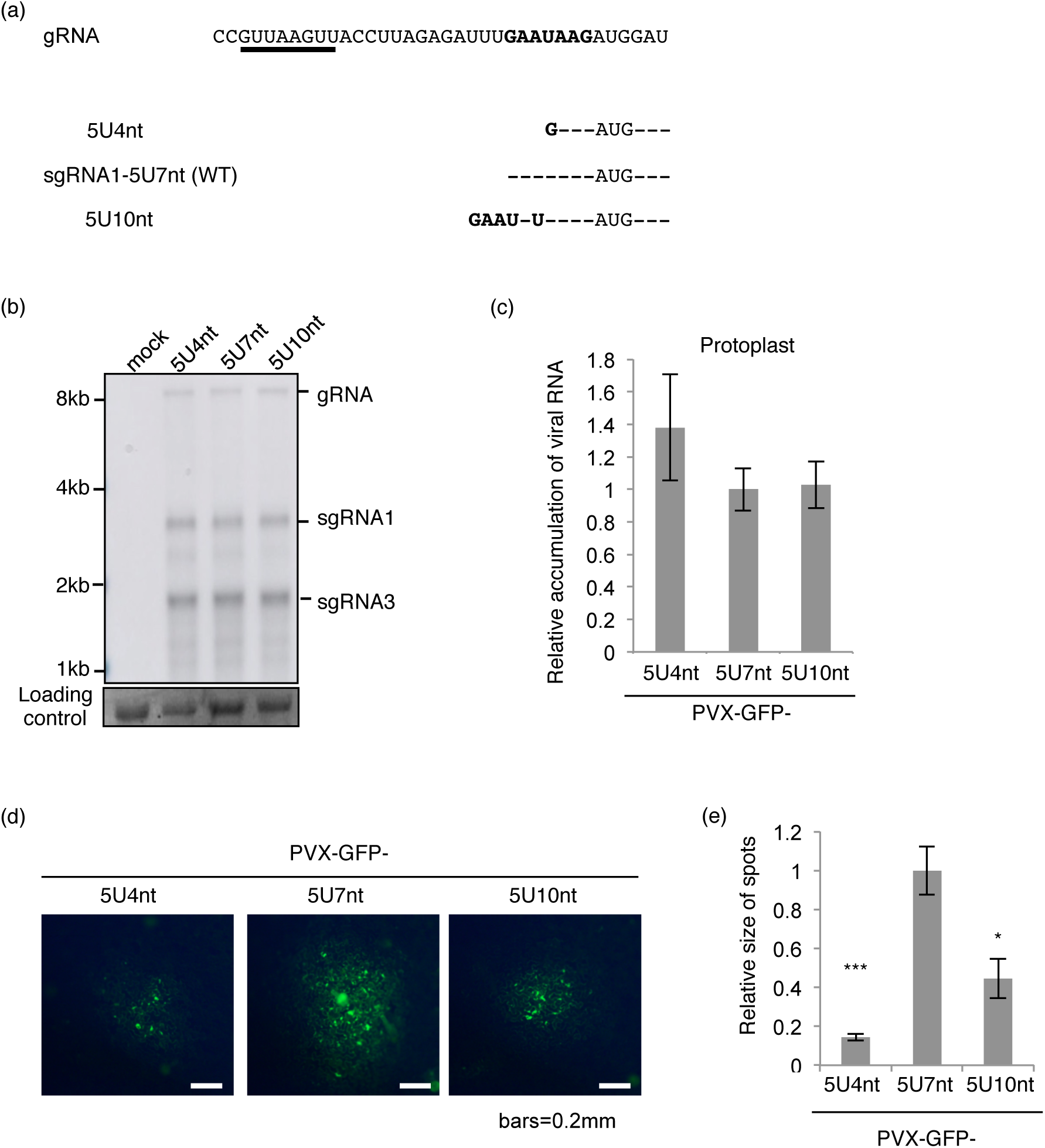
Leader-sequence length optimizes virus propagation. (a) 5′ UTR variant constructs. The putative sgRNA1 promoter sequence is underlined. Boldfaced GAAUAAG indicates the WT PVX sgRNA1 leader sequence. Hyphens indicate that the nucleotide at that position was not substituted in the WT. (b) Northern blotting of RNA extracted from *N. benthamiana*. (c) Relative viral RNA accumulation as quantified by qRT-PCR. (d) Images of the spread of GFP signals produced by PVX-GFP. (e) Quantification of the size of the fluorescent foci in inoculated leaves. Fluorescence images with more than 10 foci in (d) were processed using ImageJ v1.40 software (NIH) to measure the sizes of viral infection foci. The sizes were normalized to those of 5U7nt (WT). Asterisks indicate a significant difference compared with WT (Student’s t-test, single asterisk; P < 0.05, triple asterisk; P < 0.001).

### A short 5′ UTR is a widely conserved feature among uORFs in host plant mRNAs

Our results suggest that LISH finely controls efficiency of the translation from viral polycistronic RNAs. On the other hand, accumulating evidence has shown that the translation of a large number of eukaryotic polycistronic mRNAs is regulated by leaky scanning of the uORF AUG codon (6). Therefore, we hypothesized that LISH may be widely employed for the translational regulation of such eukaryotic polycistronic mRNAs.

In the *Arabidopsis* genome, uORFs can be found in almost 35% of mRNAs, and they are suggested to play crucial roles in controlling the translation level of the main ORFs (mORFs) via non-canonical translation mechanisms, including primarily leaky scanning. To verify the hypothesis, we computationally analyzed the positions of the 5′**-**proximal first AUGs (fAUGs) and compared them to annotated mORF AUGs (mAUGs) in the registered *Arabidopsis* mRNA database (TAIR9). Among 48,034 registered mRNAs, 21,546 mRNAs (∼44.9%) were canonical in structure with no uORF and matching fAUGs and mAUGs. Among these 21,546 canonical fAUG-mAUG-matched mRNAs (matched-mRNAs), the mAUGs of 5,339 mRNAs were located at the 5′ terminus, which was likely due to inadequate sequence analysis of the mRNA structures, and therefore, we omitted these mRNAs from further analysis. As a result, 16,207 mRNAs were regarded as canonical matched-mRNAs, whereas 26,488 were regarded as fAUG-mAUG-mismatched mRNAs (mismatched-mRNAs) that contained uORFs.

We analyzed the lengths of the 5 ′ UTRs of fAUGs in all matched- and mismatched-mRNAs. In mismatched-mRNAs, the two most frequent lengths of 5′ UTRs of fAUGs were shorter than 20 nt. Of the total 26,488 mismatched-mRNAs, 7,705 (29.1%) mismatched-mRNAs with fAUGs had 5′ UTRs shorter than 20 nt (Fig. 7). By contrast, of the total 16,207 matched-mRNAs, 1,101 (6.8%) matched-mRNAs with fAUGs had 5′ UTRs shorter than 20 nt. Interestingly, the most frequent 5′ UTR lengths in matched-mRNAs were found between 50 and 100 nt, suggesting there is some bias to avoid short UTR. Although we cannot rule out the possibility that a portion of the sequences of mismatched-mRNAs with short 5′ UTRs as well as matched-mRNAs has not been fully analyzed, these results suggested that LISH is an intrinsic mechanism of uORF-mediated translational gene regulation in plants.

**Figure 7.**
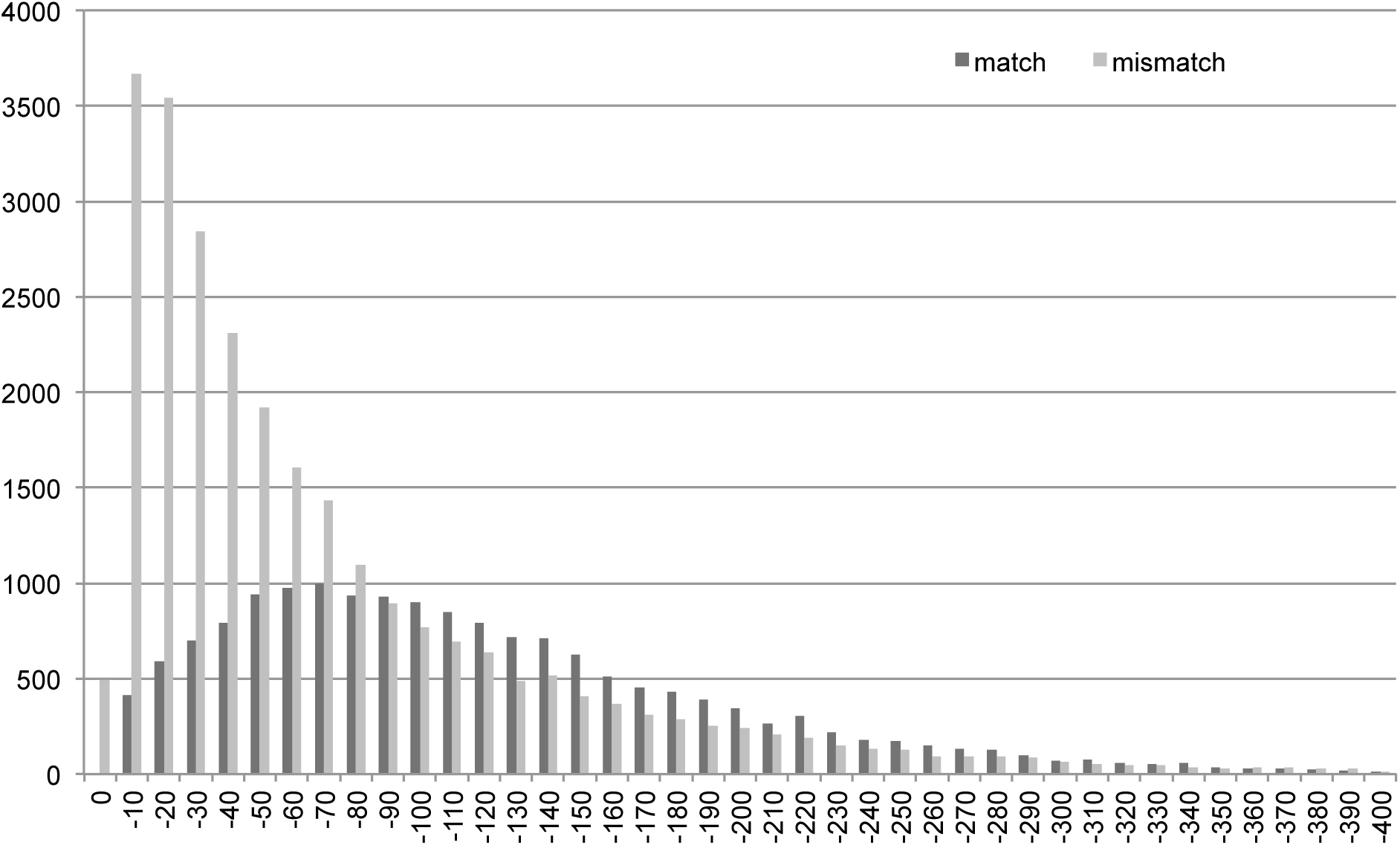
Short leader-sequence length is conserved among endogenous *Arabidopsis* uORFs. We omitted the value of matched-0 because the value was thought to be based on inadequate sequence data. The values for mRNAs whose leaders were longer than 401 nt are not shown in this graph because the values were small.

## Discussion

In this study, we elucidated a novel regulatory mechanism involving leaky scanning. We found that the TGBps encoded in the genome of potexviruses are all translated from a single RNA molecule, sgRNA1, via leaky scanning. Moreover, the remarkable shortness of the leader sequence upstream of the first ORF in sgRNA1, TGBp1, is well conserved among potexviruses. Mutational analyses showed that the efficiency of leaky scanning was strictly dependent on the 5′ UTR length. Surprisingly, genome-wide analysis of *A. thaliana* mRNAs showed that the distribution of 5′ UTR lengths in mRNAs with a uORF differed drastically from that of mRNAs lacking a uORF, indicating that 5′ UTR length-dependent regulation of leaky scanning would be another regulatory mechanism employed by not only viruses but also eukaryotes. Taken together, we present here a mechanism for the regulation of leaky scanning in addition to the Kozak sequence.

### All TGB proteins are translated from sgRNA1

We analyzed the molecular mechanism underlying the translation of three TGBps from sgRNA to re-verify the potexviral TGBps expression model proposed previously (13). In the conventional model, TGBp1 is translated from sgRNA1, whereas TGBp2 and TGBp3 are translated from sgRNA2. Our study demonstrated that in an sgRNA1-deleted PlAMV mutant (Δsg1), the accumulation of TGBp2/3 as well as TGBp1 was impaired, despite comparable level of RdRp accumulation to that in the WT. Consistent with this result, TGBp2/3 together with TGBp1 accumulated significantly in protoplasts transfected with only sgRNA1. These results indicate that TGBp2 and TGBp3 are substantially translated from sgRNA1. Furthermore, we did not find a clearly defined TSS for sgRNA2, in contrast to clearly defined TSSs for sgRNA1 and sgRNA3. It is true that there are several previous studies that have detected sgRNA2-like molecules from viruses belonging to genus *Potexvirus* (40, 41), and the possibility that these viral RNAs have some function cannot be completely ruled out. Some of the degradation intermediates of viral RNAs accumulate in large amounts and function as non-coding RNAs (42), and sgRNA2-like molecules may also accumulate under certain conditions and have some function. However, the present results indicate that sgRNA1, not sgRNA2, is responsible for the conserved function as the major template of the translation of TGBp2 and TGBp3. Our experimental results also suggest that this TGBp translation model is conserved among *Alphaflexiviridae* and *Betaflexiviridae*. Therefore, our novel model of TGBp translation, in which all TGBps are translated from sgRNA1, may be employed across a range of plant viruses including potexviruses and other related viruses.

### 5’ UTR lengths in eukaryotic mRNA, viral gRNA, and viral sgRNA

We found that the 5′ UTR of the flexiviral sgRNA1 is very short, ranging from 1 to 8 nt. Considering the 5′ UTR of flexiviral RdRp, which is translated directly from gRNA, is between 70 and 110 nt, the 5′ UTR of sgRNA1 is extremely short. Although plant virus MPs are in many cases translated from sgRNAs, the 5′ UTR of sgRNA1 of flexiviruses is much shorter than those of MP-encoding sgRNAs of other RNA viruses. For instance, the 5′ UTR length of the sgRNA encoding the MP of tobacco mosaic virus (TMV) is approximately 60 nt, which is much longer than that in flexiviruses, indicating that LISH is not required for the translation of TMV MP in harmony with its single ORF character. On the other hand, TGBp-type MPs are also encoded in several species in the genus *Benyvirus* and family *Virgaviridae*, but the 5′ UTRs of these MPs are all more than 100 nt long. This may be because TGBp2 in these viruses have been reported to be translated from its own sgRNA, not by leaky scanning of TGBp1.

Strikingly, we found that short 5′ UTRs are also conserved in plant mRNAs besides viral RNAs. The remarkable shortness of the 5′ UTR is more conserved in a large proportion of uORFs of endogenous polycistronic mRNAs in *Arabidopsis* compared to monocistronic mRNAs. Only 6.8% of the mORFs had a 5′ UTR length shorter than 20 nt, compared to 29.1% of the uORFs. Taken together, LISH may be a general translational regulatory mechanism involving leaky scanning derived from the intrinsic function of eukaryotic ribosomes, together with Kozak sequences.

### Molecular mechanism underlying LISH

In this study, we demonstrated that the efficiency of leaky scanning is regulated by the length of the 5′ UTR of uORF. Although a previous study using an *in vitro* translation system and artificial mRNA showed that it is theoretically possible that the short 5′ UTR upregulates the efficiency of leaky scanning (43, 44), this is the first study to show that LISH is actually employed to regulate the efficiency of leaky scanning in viral and eukaryotic mRNAs in living cells. Our results raise a question about the molecular mechanism by which such short mRNA leader sequences enhance the efficiency of leaky scanning. A previous study reported that when the P-site of the mammalian 43S PIC binds to the AUG initiation codon, the PIC complex structurally contacts multiple nucleotides proximal to the AUG codon, including the upstream 17 nt and the downstream 11 nt (45). Therefore, when the 5′ UTR length is shorter than 17 nt, the absence of RNA in the space inside the PIC normally occupied by the 5′ UTR sequence may alter the PIC’s overall conformation, disrupt the efficiency of start codon recognition, and thereby increase the efficiency of leaky scanning. On the other hand, given that the efficiency of leaky scanning of sgRNA1 was also dependent on the Kozak sequence even when the 5′ UTR was 7 nt (Fig. 3), 43S PIC scanning seems to still function even under the LISH condition.

### Biological significance of LISH in viral infection

Using a *trans*-complementation assay, we showed that the infection efficiency of sgRNA1-deficient PVX was optimal when sgRNA1 with a WT 5′ UTR length was complemented. The infection efficiency decreased upon complementation with sgRNA1s containing elongated or shortened 5′ UTRs (S. Fig. 6). The same result was obtained when the intact genomic sequence of a PVX-GFP infectious clone was manipulated to generate virus mutants with elongated or shortened 5′ UTRs (Fig. 6). These results suggest that the length of the sgRNA1 leader sequence is optimal for regulating the efficiency of cell-to-cell movement, potentially through fine-tuning the accumulation levels of the three TGBps. In the region upstream of the TGBp3 initiation codon in PlAMV sgRNA1, there are only two AUG codons including those of TGBp1 and TGBp2, implying evolutionary pressure to avoid unnecessary dAUGs that would impair the translation of TGBp2/3.

TGBps act as MPs, but their detailed characteristics differ slightly (46). TGBp1 is a multifunctional protein with RNA-binding and ATPase activities that acts as an MP and an RNA-silencing suppressor. TGBp1 modifies the plasmodesmata size exclusion limit and reorganizes the viral genome, proteins, host actin, and endomembranes to form the viral X body for cell-to-cell movement (46). Both TGBp2 and TGBp3 are transmembrane proteins, but they localize to different subdomains of the endoplasmic reticulum (47). Therefore, viruses may orchestrate the different functions of TGBps to adapt to the cellular environment of the host by optimizing the levels of the three MPs. In fact, the Kozak sequence of the AUGs of TGBp2 and TGBp3 are not optimal, and recognition of the TGBp1 AUG is restricted by LISH. Thus, all three TGBp accumulation is suppressed to a certain degree by LISH together with Kozak sequence regulating their translation initiation efficiency. This may be due to the fact that the accumulation of TGBps can be stressful to plant cells. PVX TGBp1 triggers cell death resulting from endoplasmic reticulum stress in *N. benthamiana* (48). Moreover, potexviral TGBp3 drastically modifies the membrane structure of the endoplasmic reticulum (49), and its overexpression causes veinal necrosis (50). This is in agreement with the finding that TGBp3 of shallot virus X and lily virus X, both are the members of genus *Potexvirus*, generally has a non-AUG initiation codon with lower translation initiation efficiency compared to the normal AUG codon (51, 52). Furthermore, a simulation study clearly demonstrated that excessive cell-to-cell movement efficiency impairs optimal selection against defective genomes and deteriorates the quality of viral genomes (39). The highest efficiency of movement does not always maximize viral fitness. Suppression of MP translation may also increase viral fitness by suppressing movement efficiency.

In conclusion, our study elucidated a novel translation regulatory mechanism involving leaky scanning, LISH, which regulates the translation efficiency of the dORF dependent on the length of the 5′ UTR of the uORF. Future study will unveil the functional universality of LISH by revealing the biological significance of the translational regulation of dORFs by LISH in not only viral RNAs but also eukaryotic mRNAs.

## Figure Legends

**Supplementary Figure 1. Analysis of the PlAMV sgRNA TSS and construction of Δsg1 mutant.**

(a) The sgRNA1 TSS was uniquely determined at G4224 (black arrow). The putative promoter sequences are underlined. The arrow indicates the TGBp1 ORF. Boldfaced AUG indicates the TGBp1 initiation codon.

(b) The sgRNA3 TSS was uniquely determined at G5339 (black arrow). The putative promoter sequences are underlined. The arrow indicates the TGBp1 ORF. Boldfaced AUG indicates the CP initiation codon.

(c) The sgRNA2 TSS was not uniquely determined. Black dots indicate the varied 5′ ends of cloned transcripts as determined by RACE analysis. There was no promoter-like sequence located in the region between the TGBp1 initiation codon and the TGBp2 initiation codon.

(d) The putative promoter sequences are underlined. U4193, G4196, and C4199 (boldfaced and indicated by black arrows in PlAMV) were substituted to C4193, A4196, and U4199, respectively (boldfaced and indicated by black arrows in PlAMV-Δsg1). Transcription of sgRNA1 starts from G4224 as shown in Supplementary Figure 1a.

**Supplementary Figure 2. Analysis of the translation of TGBp2/3 from sgRNA1.**

(a) Δstp construct. The TGBp1 stop codon (UGA, underlined) was changed to GGA, resulting in an extension of the TGBp1 ORF. Hyphens indicate that the nucleotide at that position was not changed.

(b) Immunoblot analysis of TGBp accumulation with the Δstp construct. Interestingly, modified TGBp1 with Δstp was detected mainly in the P fraction (black arrow), indicating that the C-terminal elongation caused by the mutation made the modified TGBp1 protein insoluble.

(c) TGBp2KZ variant constructs. The 5′-terminal sequence of sgRNA1 is shown. Boldfaced AUG indicates the TGBp2 initiation codon, and U at the −3 position (underlined) was changed to A, C, or G. Hyphens indicate that the nucleotide at that position was not substituted.

(d) Immunoblot analysis of the accumulation of TGBps expressed by the TGBp2KZ variants.

(e) Dual luciferase assay analysis of the translational efficiency of TGBps expressed by TGBp2KZ variants. Rluc luminescence was normalized to Fluc luminescence. Mean values ± SDs from three independent experiments are shown.

**Supplementary Figure 3. Analysis of translation starting from the second AUG of Rluc.**

(a) Rluc activity of the variants is shown. Rluc luminescence was normalized to Fluc luminescence. Mean values ± SDs from three independent experiments are shown.

(b) Rluc variant constructs. The N-terminal amino-acid sequence of Rluc is shown. The first and fourteenth amino acids (M) were changed.

(c) Translational efficiency of TGBp1 as measured using Rluc without the M14L mutation.

**Supplementary Figure 4. Translation of TGBp2/3 from sgRNA1 is dependent on leaky scanning.**

(a) Schematic of the midAUG construct. The sequence between the TGBp1 initiation codon and the TGBp2 initiation codon is shown. Boldfaced AUG indicates the initiation codon of TGBp1 (upper) and TGBp2 (lower). Six additional AUGs (black arrow) were inserted into the TGBp1 coding region via base substitution without

changing the amino-acid sequence (red box). Hyphens indicate that the nucleotide at that position was not substituted.

(b) Immunoblot analysis of the accumulation of TGBps expressed by the midAUG construct.

(c) Dual luciferase assay analysis of the translational efficiency of TGB2/3 expressed by the midAUG construct. Rluc luminescence was normalized to Fluc luminescence. Mean values ± SDs from three independent experiments are shown. Asterisk indicates a significant difference compared with WT (Student’s t-test, P < 0.05).

**Supplementary Figure 5. Leader-sequence length affects the efficiency of leaky scanning.**

(a) Construct schematic. The 5′-terminal sequence of sgRNA1 is shown. Boldfaced AUG indicates the TGBp1 initiation codon, and the leader sequence was modified.

(b) Immunoblot analysis of the accumulation of TGBps expressed by leader-length variants.

(c) Dual luciferase assay analysis of the translational efficiency of TGBps expressed by leader-length variants. Rluc luminescence was normalized to Fluc luminescence. Mean values ± SDs from three independent experiments are shown. Asterisks indicate a significant difference compared with WT (Student’s t-test, P < 0.05).

**Supplementary Figure 6. Leader-sequence length optimizes virus propagation.**

(a) 5′ UTR variant constructs. Boldfaced GAAUAAG indicates the WT PVX sgRNA1 leader sequence. Hyphens indicate that the nucleotide at that position was not changed from the WT. AUG (underlined) is the PVX TGBp1 initiation codon.

(b) Images of the spread of PVX-GFP.

(c) Quantification of the sizes of fluorescent foci in inoculated leaves. Fluorescence images with more than 10 foci in (b) were processed using ImageJ v1.40 software (NIH) to measure the size of viral infection foci. The sizes were normalized to those of 5U7nt (WT).

Supplementary Figure 7. Leader-sequence length optimizes virus propagation.

(a) 5′ UTR variant constructs. Boldfaced GAUUAAC indicates the WT PlAMV sgRNA1 leader sequence. Hyphens indicate that the nucleotide at that position was not changed from the WT. AUG (underlined) is the PlAMV TGBp1 initiation codon.

(b) Images of the spread of PlAMV-GFP.

(c) Quantification of the sizes of fluorescent foci in inoculated leaves. Fluorescence images with more than 10 foci in (b) were processed using ImageJ v1.40 software (NIH) to measure the sizes of viral infection foci. The sizes were normalized to those of 5U7nt (WT). Asterisks indicate a significant difference compared with WT (Student’s t-test, single asterisk; P < 0.05, triple asterisk; P < 0.001).

