## supplementary information for "Short 5′ UTR enables optimal translation of plant virus tricistronic RNA via leaky scanning"

### Supplementary Figure 1. **Analysis of the PlAMV sgRNA TSS.**

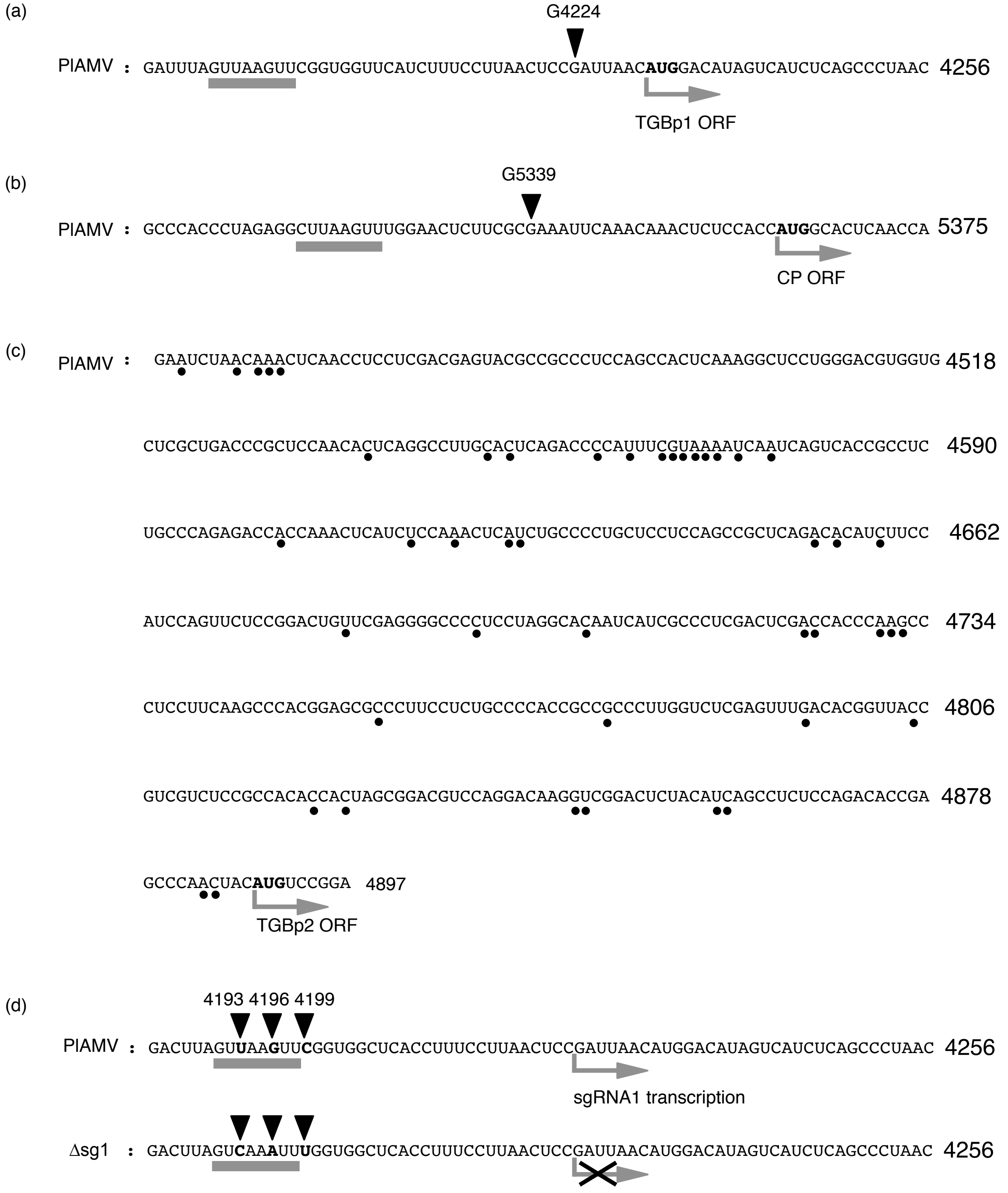


Supplementary Figure 1. **Analysis of the PlAMV sgRNA TSS and construction of PlAMV-Δsg1 mutant.**

(a) The sgRNA1 TSS was uniquely determined at G4224 (black arrow). The putative promoter sequences are underlined. The arrow indicates the TGBp1 ORF. Boldfaced AUG indicates the TGBp1 initiation codon.

(b) The sgRNA3 TSS was uniquely determined at G5339 (black arrow). The putative promoter sequences are underlined. The arrow indicates the TGBp1 ORF. Boldfaced AUG indicates the CP initiation codon.

(c) The sgRNA2 TSS was not uniquely determined. Black dots indicate the varied 5' ends of cloned transcripts as determined by RACE analysis. There was no promoter-like sequence located in the region between the TGBp1 initiation codon and the TGBp2 initiation codon.

(d) Schematic of PlAMV-Δsg1 constructs. The putative promoter sequences are underlined. U4193, G4196, and C4199 (boldfaced and indicated by black arrows in PlAMV) were substituted to C4193, A4196, and U4199, respectively (boldfaced and indicated by black arrows in PlAMV-Δsg1). Transcription of sgRNA1 starts from G4224 as shown in Supplementary Figure 1a.

Supplementary Figure 2. **Analysis of the translation of TGBp2/3 from sgRNA1.**

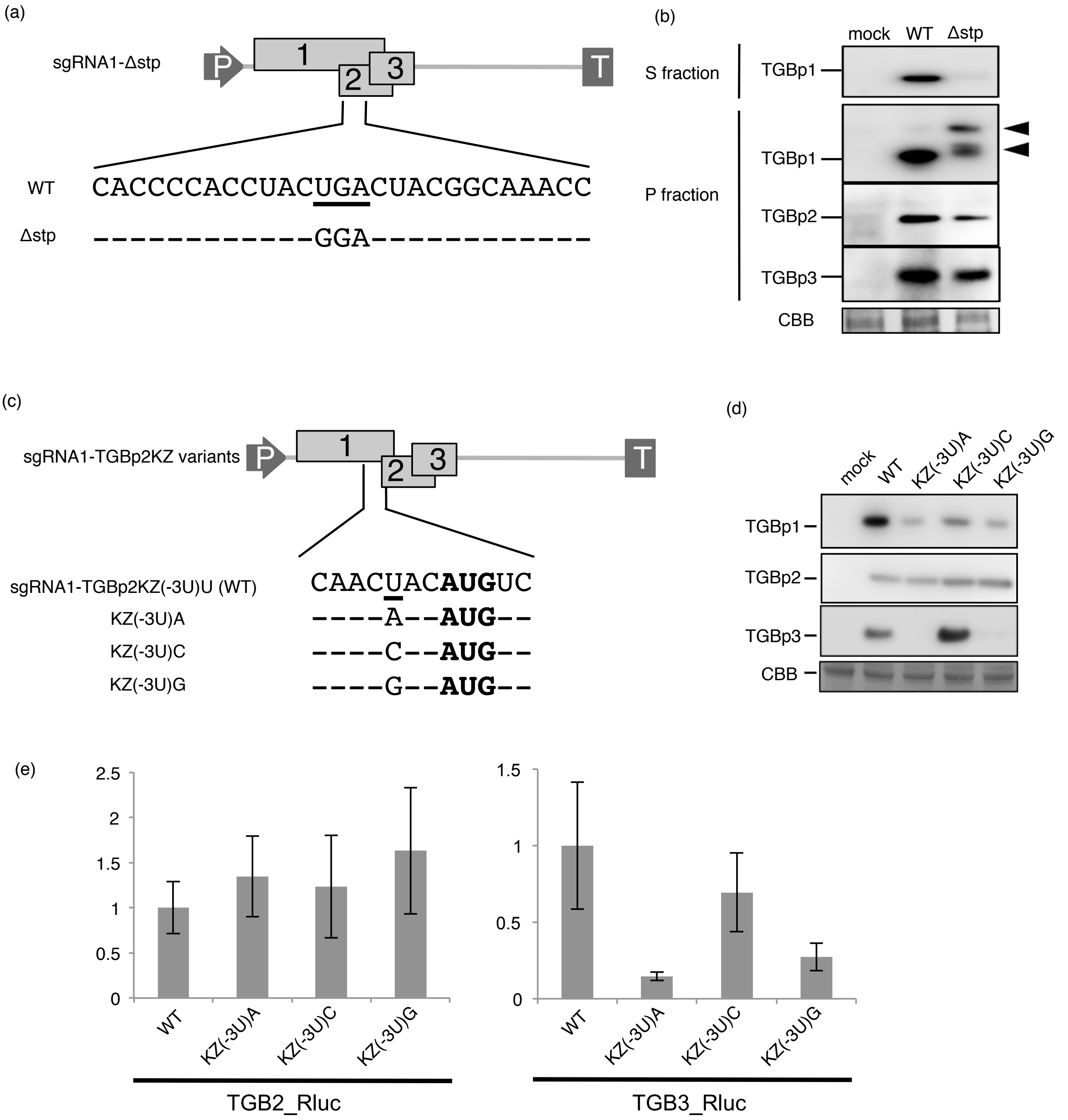


Supplementary Figure 2. **Analysis of the translation of TGBp2/3 from sgRNA1.**

(a) Δstp construct. The TGBp1 stop codon (UGA, underlined) was changed to GGA, resulting in an extension of the TGBp1 ORF. Hyphens indicate that the nucleotide at that position was not changed.

(b) Immunoblot analysis of TGBp accumulation with the Δstp construct. Interestingly, modified TGBp1 with Δstp was detected mainly in the P fraction (black arrow), indicating that the C-terminal elongation caused by the mutation made the modified TGBp1 protein insoluble.

(c) TGBp2KZ variant constructs. The 5'-terminal sequence of sgRNA1 is shown. Boldfaced AUG indicates the TGBp2 initiation codon, and U at the -3 position (underlined) was changed to A, C, or G. Hyphens indicate that the nucleotide at that position was not substituted.

(d) Immunoblot analysis of the accumulation of TGBps expressed by the TGBp2KZ variants.

(e) Dual luciferase assay analysis of the translational efficiency of TGBps expressed by TGBp2KZ variants. Rluc luminescence was normalized to Fluc luminescence. Mean values ± SDs from three independent experiments are shown.

Supplementary Figure 3. **Analysis of translation starting from the second AUG of Rluc.**

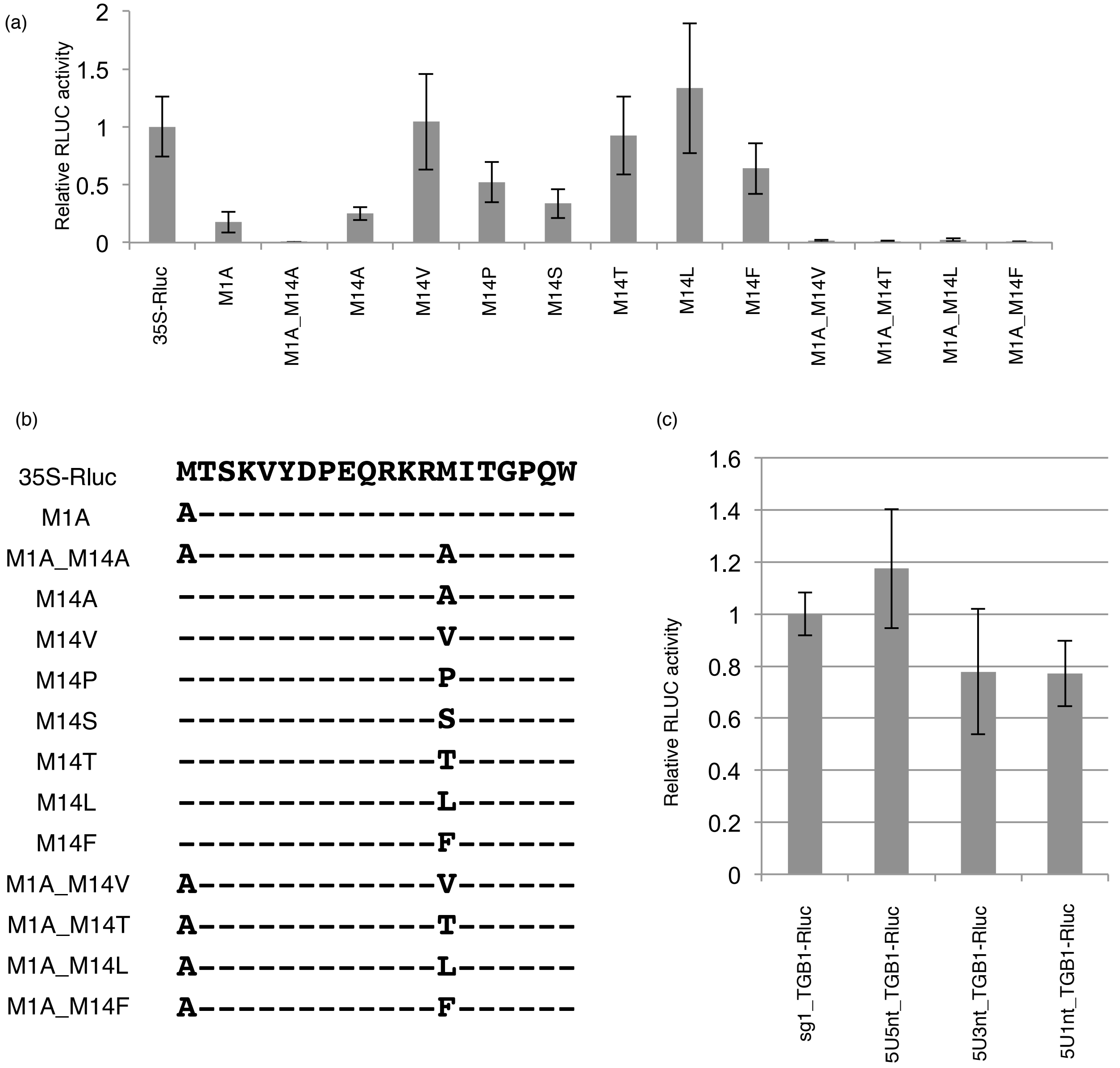


Supplementary Figure 3. **Analysis of translation starting from the second AUG of Rluc.**

(a) Rluc activity of the variants is shown. Rluc luminescence was normalized to Fluc luminescence. Mean values ± SDs from three independent experiments are shown.

(b) Rluc variant constructs. The N-terminal amino-acid sequence of Rluc is shown. The first and fourteenth amino acids (M) were changed.

(c) Translational efficiency of TGBp1 as measured using Rluc without the M14L mutation.

Supplementary Figure 4. **Translation of TGBp2/3 from sgRNA1 is dependent on leaky scanning.**

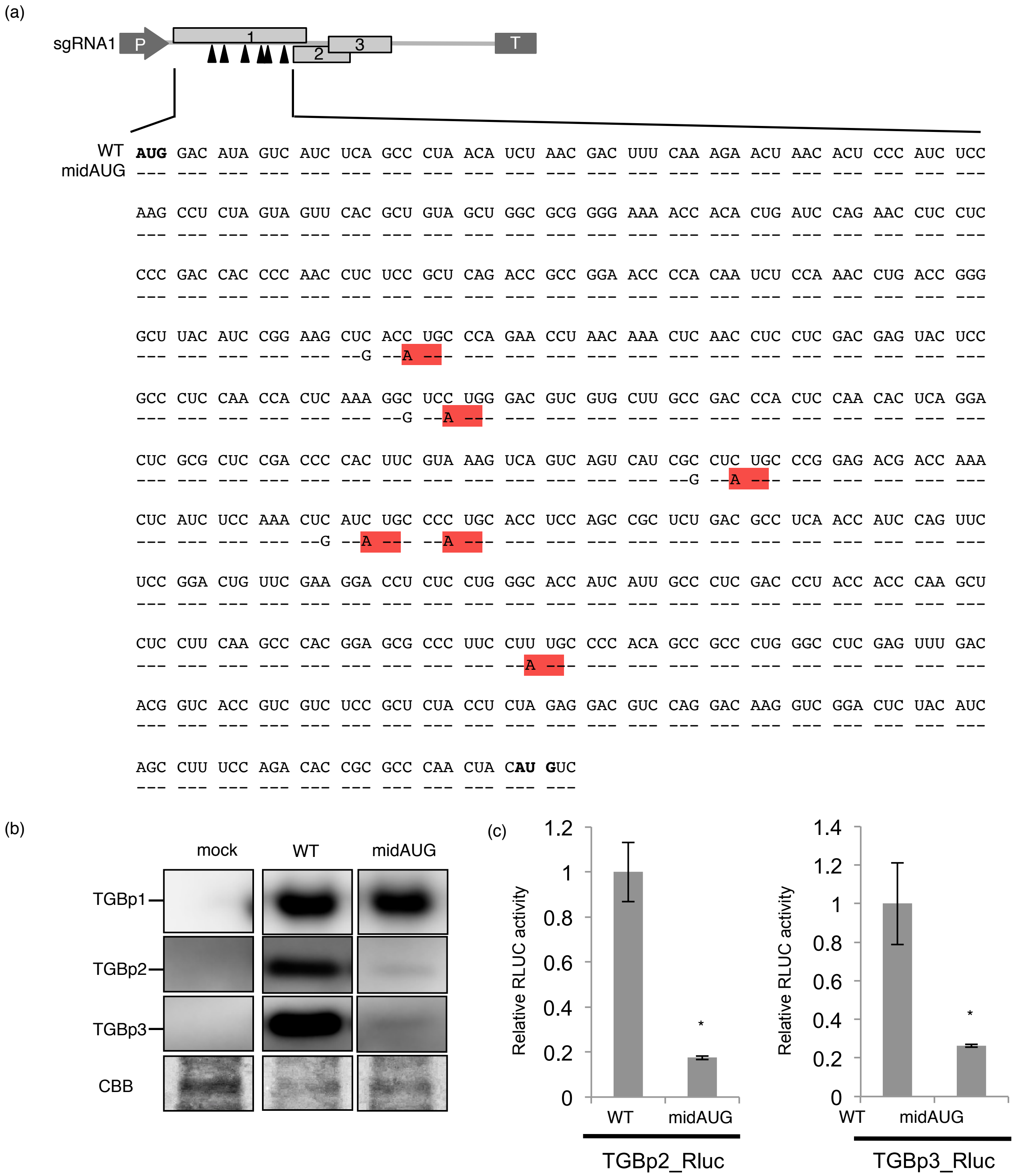


Supplementary Figure 4. **Translation of TGBp2/3 from sgRNA1 is dependent on leaky scanning.**

(a) Schematic of the midAUG construct. The sequence between the TGBp1 initiation codon and the TGBp2 initiation codon is shown. Boldfaced AUG indicates the initiation codon of TGBp1 (upper) and TGBp2 (lower). Six additional AUGs (black arrow) were inserted into the TGBp1 coding region via base substitution without changing the amino-acid sequence (red box). Hyphens indicate that the nucleotide at that position was not substituted.

(b) Immunoblot analysis of the accumulation of TGBps expressed by the midAUG construct.

(c) Dual luciferase assay analysis of the translational efficiency of TGB2/3 expressed by the midAUG construct. Rluc luminescence was normalized to Fluc luminescence. Mean values ± SDs from three independent experiments are shown. Asterisk indicates a significant difference compared with WT (Student's t-test, $P < 0.05$).

Supplementary Figure 5. **Leader-sequence length affects the efficiency of leaky scanning.**

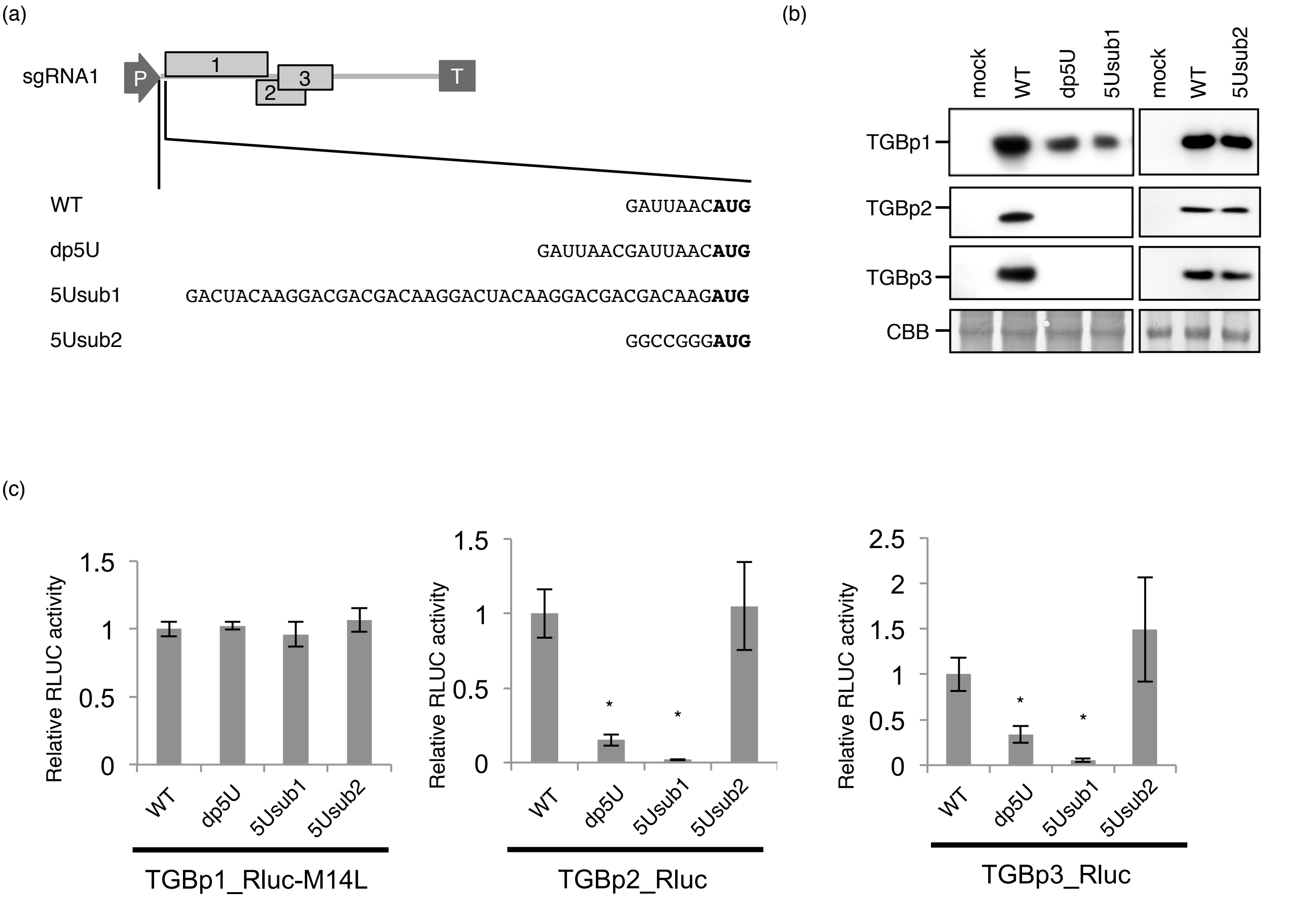


Supplementary Figure 5. **Leader-sequence length affects the efficiency of leaky scanning.**

(a) Construct schematic. The 5'-terminal sequence of sgRNA1 is shown. Boldfaced AUG indicates the TGBp1 initiation codon, and the leader sequence was modified.

(b) Immunoblot analysis of the accumulation of TGBps expressed by leader-length variants.

(c) Dual luciferase assay analysis of the translational efficiency of TGBps expressed by leader-length variants. Rluc luminescence was normalized to Fluc luminescence. Mean values ± SDs from three independent experiments are shown. Asterisks indicate a significant difference compared with WT (Student’s t-test, $P < 0.05$).

Supplementary Figure 6. **Leader-sequence length optimizes virus propagation.**

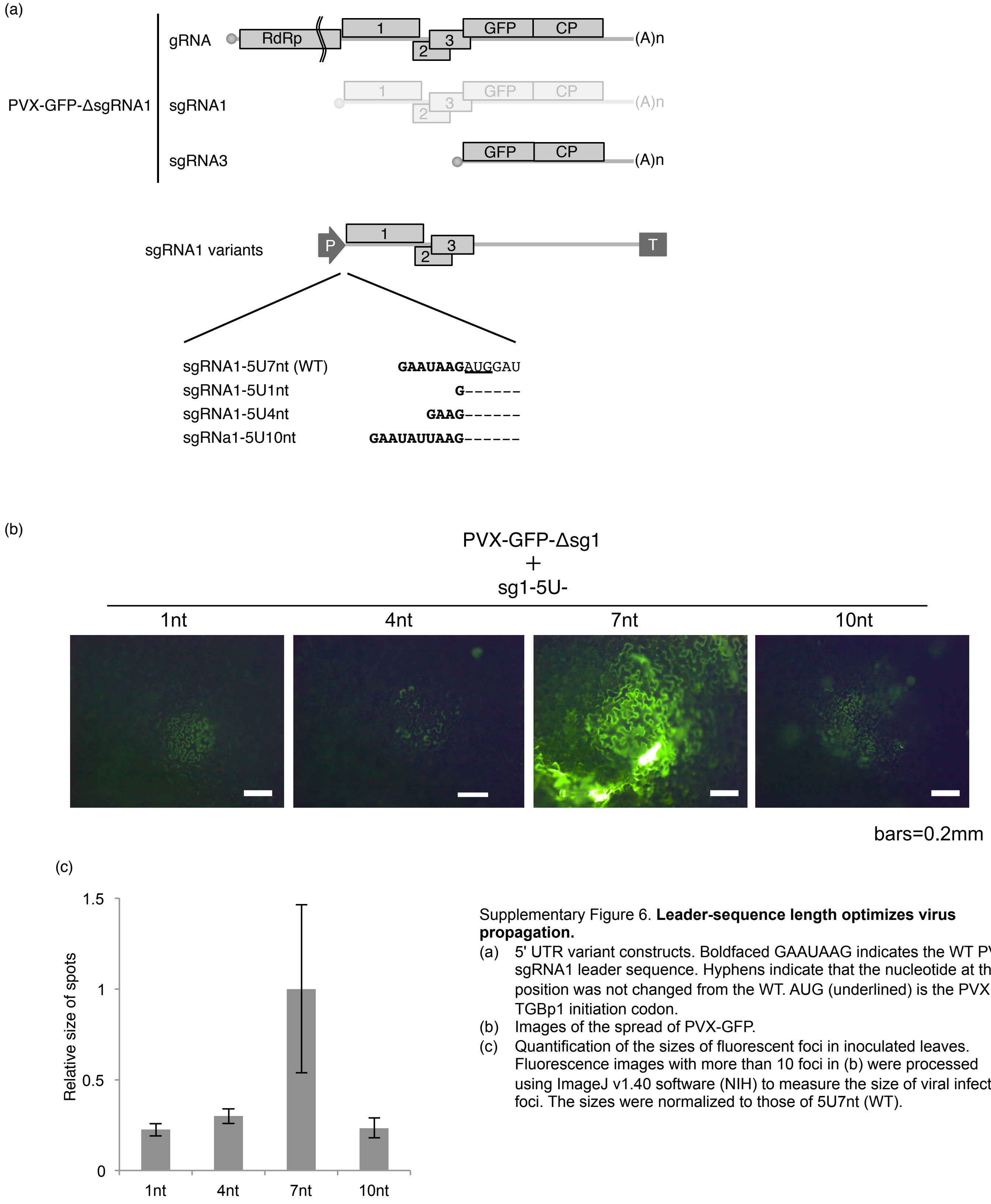


Supplementary Figure 6. **Leader-sequence length optimizes virus propagation.**

(a) 5' UTR variant constructs. Boldfaced GAAUAAG indicates the WT PVX sgRNA1 leader sequence. Hyphens indicate that the nucleotide at that position was not changed from the WT. AUG (underlined) is the PVX TGBp1 initiation codon.

(b) Images of the spread of PVX-GFP.

(c) Quantification of the sizes of fluorescent foci in inoculated leaves. Fluorescence images with more than 10 foci in (b) were processed using ImageJ v1.40 software (NIH) to measure the size of viral infection foci. The sizes were normalized to those of 5U7nt (WT).

Supplementary Figure 7. **Leader-sequence length optimizes virus propagation.**

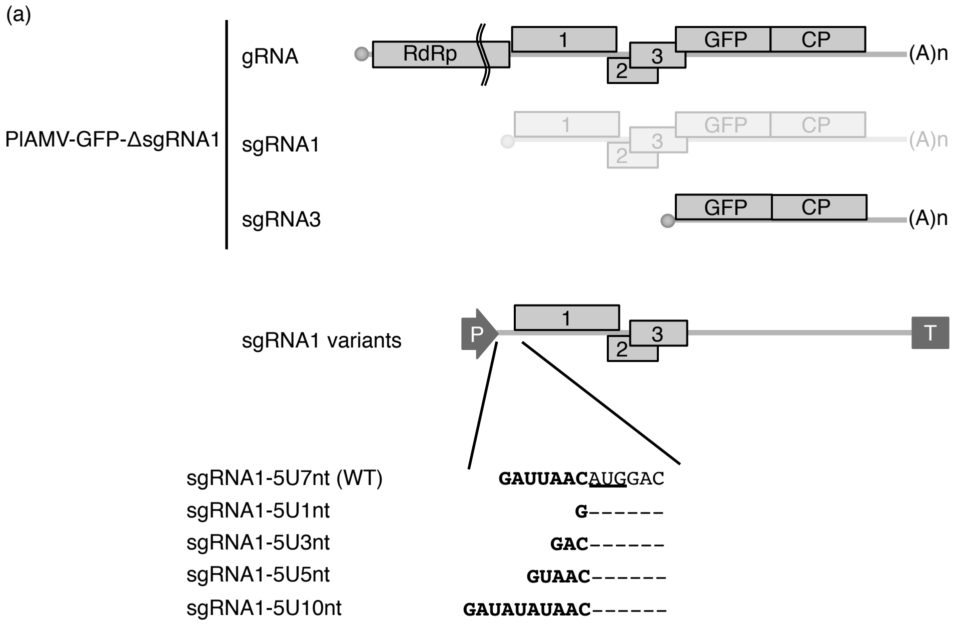


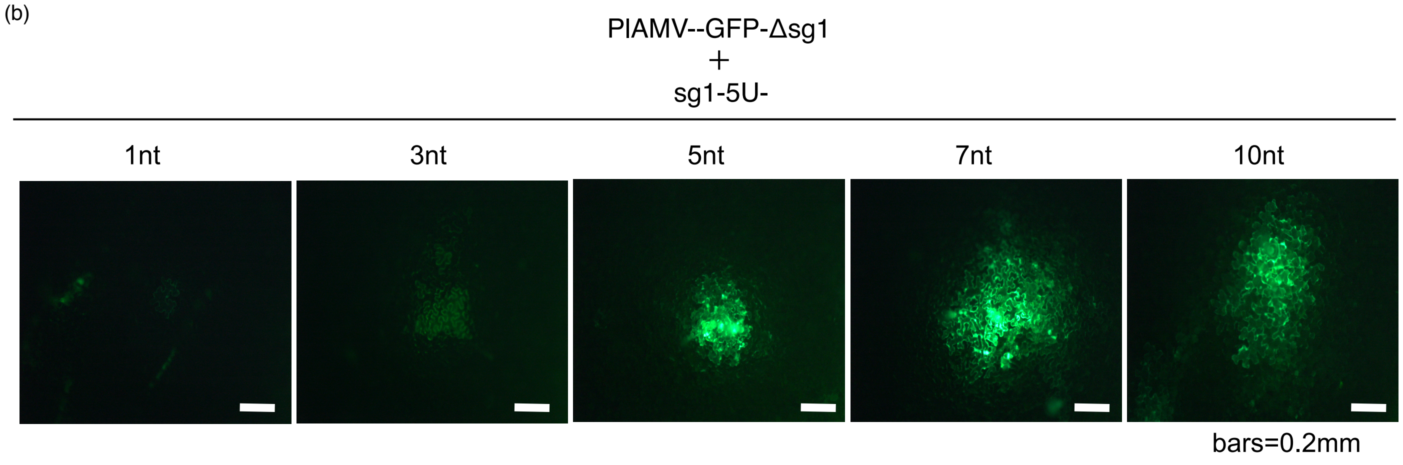


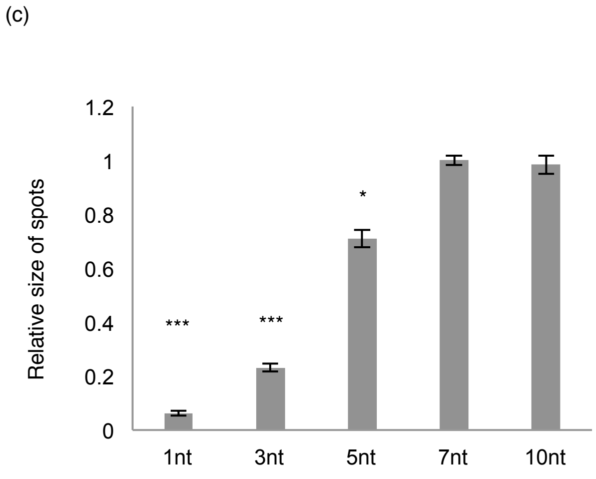


Supplementary Figure 7. **Leader-sequence length optimizes virus propagation.**

(a) 5' UTR variant constructs. Boldfaced GAUUAAC indicates the WT PlAMV sgRNA1 leader sequence. Hyphens indicate that the nucleotide at that position was not changed from the WT. AUG (underlined) is the PlAMV TGBp1 initiation codon.

(b) Images of the spread of PlAMV-GFP.

(c) Quantification of the sizes of fluorescent foci in inoculated leaves. Fluorescence images with more than 10 foci in (b) were processed using ImageJ v1.40 software (NIH) to measure the sizes of viral infection foci. The sizes were normalized to those of 5U7nt (WT). Asterisks indicate a significant difference compared with WT (Student's t-test, single asterisk; $P < 0.05$, triple asterisk; $P < 0.001$).

### Supplementary Table 1

| target | sequence |
|---|---|
| PVX-TGBp1 | full length |
| PVX-TGBp2 | CGNNHSSH+SLPHGGAYRDGTK |
| LoLV-TGBp1 | GFERTDEPLPSDS |
| LoLV-TGBp2 | ADPHHPTV |
| LoLV-CP | full length |
| PVM-TGBp1 | LHKYKFERLNNKLA |
| PVM-TGBp2 | GDRDHRLPHGGWY |
| PVM-CP | AKEAGTSQAAKGNR |

Supplementary Table 2

| midAUGinsert |
| --- |
| GGAAGTTCATTTCATTTGGAGAGGACGATTAACATGGACATAGTCATCTCAGCCCTAACATCTAACGACTTTCAAAGAACTAACACTCCCATCTCCAAGCCTCTAGTAGT<br>TCACGCTGTAGCTGGCGCGGGGAAAACCACACTGATCCAGAACCTCCTCCCCGACCACCCCAACCTCTCCGCTCAGACCGCCGGAACCCCACAATCTCCAAACCTGAC<br>CGGGGCTTACATCCGGAAGCTGACATGCCCAGAACCTAACAAACTCAACCTCCTCGACGAGTACTCCGCCCTCCAACCACTCAAAGGGTCATGGGACGTCGTGCTTGC<br>CGACCCACTCCAACACTCAGGACTCGCGCTCCGACCCCACTTCGTAAAGTCAGTCAGTCATCGGCTATGCCCGGAGACGACCAAACTCATCTCCAAACTGATATGCCCA<br>TGCACCTCCAGCCGCTCTGACGCCTCAACCATCCAGTTCTCCGGACTGTTCGAAGGACCTCTCCTGGGCACCATCATTGCCCTCGACCCTACCACCCAAGCTCTCCTTCA<br>AGCCCACGGAGCGCCCTTCCTATGCCCCACAGCCGCCCTGGG |

Supplementary Table 3

| Primer name | Primer sequence |
|---|---|
| PepMV1F | GAAAACAAAACATAACACATAATATC |
| GRR | GGGGTACCGCTACGTAACGGCATGACAGTGTTTTTTTTTTTTTTTTTTTTTTTT |
| PVM1F | GATAAACAAACATACAATATCTGGACTTACACTGC |
| GRF | CCGTTACGTAGCGGTACCCCTCAAACATTTGGCAATAAA |
| PVM35SR | ATATTGTATGTTTGTTTATCCCTCTCCAAATGAAATGAAC |
| PepMV35SR | ATGTGTTATGTTTTGTTTTCCCTCTCCAAATGAAATGAAC |
| 35SF | TCGGGAAACCTCCTCGGATTCCATTGCCCAGC |
| 35SR | AATGGAATCCGAGGAGGTTTCCCGATATTACCC |
| NOSF | ATGATTAGAGTCCCGCAATTATACATTTAATACGC |
| NOSR | TGTATAATTGCGGGACTCTAATCATAAAAACCC |
| PlAMVdsg1F | CGGTTCGTGACTTAGTCAAATTTGGTGGCTCACCTTTCCTTAAC |
| PlAMVdsg1R | AAATTTGACTAAGTCACGAACCGTTACC |
| LoLVdsg1F | GTCTTTTGGTTTTTGGTTACCATACCCACTAGTCGATTAAC |
| LoLVdsg1R | GTTAATCGACTAGTGGGTATGGTAACCAAAAACCAAAAGAC |
| PVMdsg1F | GTTAGTGTAGCTTACCAAGTGCTATTGTATTGAATATTTATGG |
| PVMdsg1R | ATAGCACTTGGTAAGCTACACTAACCAATCAGAGC |
| OSdsg1F | TACCTTATAGATTTGAATAAGATGGATATTCTC |
| OSdsg1R | TGATAACGGTTAAAGAAAGTTTCTGAGGC |
| 35S-fluc-1F | GACTCTAGAGGATCCATGGAAGACGCCAAAAACATAAAGAAAGGC |
| 35S-fluc-1653R | AGCCGGGCGGCCGCTTTACACGGCGATCTTTCCGCCCTTC |
| 35S-down1F | AGCGGCCGCCCGGCTGCAGATCG |
| 35S-up1R | GGATCCTCTAGAGTCGACTG |
| M1AF | GCGACTTCGAAAGTTTATGATCCAGAAC |
| M14AF | GCGATAACTGGTCCGCAGTGGTGG |
| M14FF | TTCATAACTGGTCCGCAGTGGTGG |
| M14LF | CTTATAACTGGTCCGCAGTGGTGG |
| M14PF | CCAATAACTGGTCCGCAGTGGTGG |
| M14SF | AGCATAACTGGTCCGCAGTGGTGG |
| M14TF | ACTATAACTGGTCCGCAGTGGTGG |
| M14VF | GTTATAACTGGTCCGCAGTGGTGG |
| M14R | CCGTTTCCTTTGTTCTGGATC |
| 35S-4224F | TCATTTGGAGAGGACGATTAACATGGACATAGTCATCTCAGCCC |
| 35S-6102R | AGCCGGGCGGCCGCTGGCCCACCAGACTTTCACTGGTCG |
| 35S-up96R | GTCCTCTCCAAATGAAATGAACTTCC |
| Rluc-1F | ATGACTTCGAAAGTTTATGATCCAG |
| Rluc-936R | TTATTGTTCATTTTTGAGAACTCGC |
| 35S-TGB15U-Rluc-R | GAAGTCATGTTAATCGTCCTCTCCAAATGAAATGAACTTCC |
| TGB25U-Rluc-R | AACTTTCGAAGTCATGTAGTTGGGCGCGGTGTCTGG |
| TGB35U-Rluc-R | AACTTTCGAAGTCATGAAGGTTGTCGCCGACGTGGGGGAG |
| Rluc-TGB13U-F | CAAAAATGAACAATAACTACGGCAAACCCGTTCTCGCTGCTTCC |
| Rluc-TGB23U-F | CAAAAATGAACAATAAACCCGGCTGCACCATTGTGATCACAGGG |
| Rluc-TGB33U-F | CAAAAATGAACAATAACGCCCTCAACGCGATGACCTTCG |
| KSF | GATTAACGGGGCGCGTGGTGGCGGCTGCAGCCGCCACCACGCGCCCCATGGACATAGTCATCTCAGCC |
| KSRlucF | GATTAACGGGGCGCGTGGTGGCGGCTGCAGCCGCCACCACGCGCCCCATGACTTCGAAAGTTTATGATCCAG |
| dstpF | CACCTACGGACTACGGCAAACCCGTTCTCGC |
| PlAMV-4913R | GGGTGAGGTGGTGGGCTCCGGAC |
| KZUF | GATTTACATGGACATAGTCATCTCAGCCC |
| KZCF | GATTCACATGGACATAGTCATCTCAGCCC |
| KZGF | GATTGACATGGACATAGTCATCTCAGCCC |
| KZURlucF | GATTTACATGACTTCGAAAGTTTATGATCCAG |
| KZCRlucF | GATTCACATGACTTCGAAAGTTTATGATCCAG |
| KZGRlucF | GATTGACATGACTTCGAAAGTTTATGATCCAG |
| 1ntR | CGTCCTCTCCAAATGAAATGAACTTCC |
| 3ntR | GTCGTCCTCTCCAAATGAAATGAACTTCC |
| 5ntR | GTTATCGTCCTCTCCAAATGAAATGAACTTCC |
| TGB1F | ATGGACATAGTCATCTCAGCCC |
| dpF | GATTAACGATTAACATGGACATAGTCATCTC |
| dpRlucF | GATTAACGATTAACATGACTTCGAAAGTTTATGATCCAG |
| sub1F | GACTACAAGGACGACGACAAGGACTACAAGGACGACGACAAGATGGACATAGTCATCTCAGCCC |
| sub1RlucF | GACTACAAGGACGACGACAAGGACTACAAGGACGACGACAAGATGACTTCGAAAGTTTATGATCCAG |
| sub2F | GGCCGGGATGGACATAGTCATCTCAGCCC |
| sub2RlucF | GGCCGGGATGACTTCGAAAGTTTATGATCCAG |
| TGB2ATGF | ATGTCCGGAGCCCACCACCTCAC |
| TGB2KZTAR | GTTGTTGGGCGCGGTGTCTGGAAAGG |
| TGB2KZTCR | GTGGTTGGGCGCGGTGTCTGGAAAGG |
| TGB2KZTGR | GTCGTTGGGCGCGGTGTCTGGAAAGG |
| 35S-flag-GFP1F | GGAAGTTCATTTCATTTGGAGAGGACGACTACAAGGACGACGACAAGATGGTGAGCAAGGGCGAGGAG |
| GFP-5U-TGB1R | GGGCTGAGATGACTATGTCCATGTTAATCTTACTTGTACAGCTCGTCCATGC |
| GFP-5U-RlucR | CTGGATCATAAACTTTCGAAGTCATGTTAATCTTACTTGTACAGCTCGTCCATGC |
| midAUGF | CTATGCCCCACAGCCGCCCTGGG |
| RlucF | GGGCCAGATGTAAACAAATGAATG |
| RlucR | ATTTGCCTGATTTGCCCATAC |
| FlucF | CGGAGGAGTTGTGTTTGTGG |
| FlucR | ATCTTTCCGCCCTTCTTGG |
| actin2F | GCACCCTGTTCTTCTTACCG |
| actin2R | AACCCTCGTAGATTGGCACA |
| PVX5435F | AATACATATCTCAACGCAATCATACTTG |
| PVX-RT7 | TAATACGACTCACTATAGGGATTTATATTATTCATACAATCAAACCAGAAAATACTATGAAAC |
| LoLV6611F | ATGTCAGAATCCAAAGCAGAGAC |
| LoLVRT7 | TAATACGACTCACTATAGGGGCTTTGACGGCAAAACCGAGGG |
| PVM7531F | TCGCTTGAGGCACTGAGCAG |
| PVM-RT7 | TAATACGACTCACTATAGGGAGGCTAAAAATAGTTAAAAACCTAGTTTTATTTATAGTAG |
| PrGRR | GGGGTACCGCTACGTAACGGCATGACAGTGTTTTTTTTTTTTTTTTTTTTTTTTGGCCCACCAGACTTTCACTGGTCG |
| 35S-Prsg1-5U1ntF | GTTCATTTCATTTGGAGAGGGATGGACATAGTCATCTCAGCCCT |
| 35S-Prsg1-5U3ntF | GTTCATTTCATTTGGAGAGGGACATGGACATAGTCATCTCAGCCCT |
| 35S-Prsg1-5U5ntF | GTTCATTTCATTTGGAGAGGGTAACATGGACATAGTCATCTCAGCCCT |
| 35S-Prsg1-5U7ntF | GTTCATTTCATTTGGAGAGGGATTAACATGGACATAGTCATCTCAGCCCT |
| 35S-Prsg1-5U10ntF | GTTCATTTCATTTGGAGAGGGATATATAACATGGACATAGTCATCTCAGCCCT |
| OS-TGB1F | ATGGATATTCTCATCATTAGTTTGAAAAGTTTAGG |
| OS-sg1-5U4ntR | CAAACTAATGATGAGAATATCCATCTTCAAAACTATAAGGTAACTTAACGG |
| OS-sg1-5U10ntR | TCAAACTAATGATGAGAATATCCATCTTAATATTCAAATCTATAAGGTAACTT |
| 35S-OS-5U1nt-F | TTTCATTTGGAGAGGGATGGATATTCTCATCATTAGTTTGAAAAG |
| 35S-OS-5U4nt-F | TTTCATTTGGAGAGGGAAGATGGATATTCTCATCATTAGTTTGAAAAG |
| 35S-OS-5U7nt-F | TTTCATTTGGAGAGGGAATAAGATGGATATTCTCATCATTAGTTTGAAAAG |
| 35S-OS-5U10nt-F | TTTCATTTGGAGAGGGAATATTAAGATGGATATTCTCATCATTAGTTTGAAAAG |
| OSGRR | GGGGTACCGCTACGTAACGGCATGACAGTGTTTTTTTTTTTTTTTTTTTTTTTTATTTATATTATTCATACAATCAAACCAG |

#### Supplementary Table 4

| construction name | Name | template | F | R |
|---|---|---|---|---|
| **Leader sequence variant of sgRNA1** | **Δstp** | 35S-PlAMV-sg1 | dstpF | PlAMV-4913R |
| | **KS-sg1** | 35S-PlAMV-sg1 | KSF | 35S-up96R |
| | **KZ(-3A)U** | 35S-PlAMV-sg1 | KZUF | 35S-up96R |
| | **KZ(-3A)C** | 35S-PlAMV-sg1 | KZCF | 35S-up96R |
| | **KZ(-3A)G** | 35S-PlAMV-sg1 | KZGF | 35S-up96R |
| | **5U5nt** | 35S-PlAMV-sg1 | TGB1-1F | 5ntR |
| | **5U3nt** | 35S-PlAMV-sg1 | TGB1-1F | 3ntR |
| | **5U1nt** | 35S-PlAMV-sg1 | TGB1-1F | 1ntR |
| | **dp5U** | 35S-PlAMV-sg1 | dpF | 35S-up96R |
| | **5Usub1** | 35S-PlAMV-sg1 | sub1F | 35S-up96R |
| | **5Usub2** | 35S-PlAMV-sg1 | sub2F | 35S-up96R |
| **Leader sequence variant of sgRNA1-TGBp1_Rluc** | **KS** | sgRNA1-TGBp1_Rluc | KSRlucF | 35S-up96R |
| | **KZ(-3A)U** | sgRNA1-TGBp1_Rluc_M14L | KZURlucF | 35S-up96R |
| | **KZ(-3A)C** | sgRNA1-TGBp1_Rluc_M14L | KZCRlucF | 35S-up96R |
| | **KZ(-3A)G** | sgRNA1-TGBp1_Rluc_M14L | KZGRlucF | 35S-up96R |
| | **5U5nt** | sgRNA1-TGBp1_Rluc_M14L | Rluc-1F | 5ntR |
| | **5U3nt** | sgRNA1-TGBp1_Rluc_M14L | Rluc-1F | 3ntR |
| | **5U1nt** | sgRNA1-TGBp1_Rluc_M14L | Rluc-1F | 1ntR |
| | **dp5U** | sgRNA1-TGBp1_Rluc_M14L | dpRlucF | 35S-up96R |
| | **5Usub1** | sgRNA1-TGBp1_Rluc_M14L | sub1RlucF | 35S-up96R |
| | **5Usub2** | sgRNA1-TGBp1_Rluc_M14L | sub2RlucF | 35S-up96R |
| **Leader sequence variant of sgRNA1-TGBp2_Rluc** | **KS** | sgRNA1-TGBp2_Rluc | KSF | 35S-up96R |
| | **KZ(-3A)U** | sgRNA1-TGBp2_Rluc | KZUF | 35S-up96R |
| | **KZ(-3A)C** | sgRNA1-TGBp2_Rluc | KZCF | 35S-up96R |
| | **KZ(-3A)G** | sgRNA1-TGBp2_Rluc | KZGF | 35S-up96R |
| | **5U5nt** | sgRNA1-TGBp2_Rluc | TGB1-1F | 5ntR |
| | **5U3nt** | sgRNA1-TGBp2_Rluc | TGB1-1F | 3ntR |
| | **5U1nt** | sgRNA1-TGBp2_Rluc | TGB1-1F | 1ntR |
| | **dp5U** | sgRNA1-TGBp2_Rluc | dpF | 35S-up96R |
| | **5Usub1** | sgRNA1-TGBp2_Rluc | sub1F | 35S-up96R |
| | **5Usub2** | sgRNA1-TGBp2_Rluc | sub2F | 35S-up96R |
| **Leader sequence variant of sgRNA1-TGBp3_Rluc** | **KS** | sgRNA1-TGBp3_Rluc | KSF | 35S-up96R |
| | **KZ(-3A)U** | sgRNA1-TGBp3_Rluc | KZUF | 35S-up96R |
| | **KZ(-3A)C** | sgRNA1-TGBp3_Rluc | KZCF | 35S-up96R |
| | **KZ(-3A)G** | sgRNA1-TGBp3_Rluc | KZGF | 35S-up96R |
| | **5U5nt** | sgRNA1-TGBp3_Rluc | TGB1-1F | 5ntR |
| | **5U3nt** | sgRNA1-TGBp3_Rluc | TGB1-1F | 3ntR |
| | **5U1nt** | sgRNA1-TGBp3_Rluc | TGB1-1F | 1ntR |
| | **dp5U** | sgRNA1-TGBp3_Rluc | dpF | 35S-up96R |
| | **5Usub1** | sgRNA1-TGBp3_Rluc | sub1F | 35S-up96R |
| | **5Usub2** | sgRNA1-TGBp3_Rluc | sub2F | 35S-up96R |
| **TGBp2KZ variant of sgRNA1** | **KZ(-3U)A** | 35S-PlAMV-sg1 | TGB2ATGF | TGB2KZTAR |
| | **KZ(-3U)C** | 35S-PlAMV-sg1 | TGB2ATGF | TGB2KZTCR |
| | **KZ(-3U)G** | 35S-PlAMV-sg1 | TGB2ATGF | TGB2KZTGR |
| **TGBp2KZ variant of sgRNA1-TGBp2_Rluc** | **KZ(-3U)A** | sgRNA1-TGBp2_Rluc | Rluc-1F | TGB2KZTAR |
| | **KZ(-3U)C** | sgRNA1-TGBp2_Rluc | Rluc-1F | TGB2KZTCR |
| | **KZ(-3U)G** | sgRNA1-TGBp2_Rluc | Rluc-1F | TGB2KZTGR |
| **TGBp2KZ variant of sgRNA1-TGBp3_Rluc** | **KZ(-3U)A** | sgRNA1-TGBp3_Rluc | TGB2ATGF | TGB2KZTAR |
| | **KZ(-3U)C** | sgRNA1-TGBp3_Rluc | TGB2ATGF | TGB2KZTCR |
| | **KZ(-3U)G** | sgRNA1-TGBp3_Rluc | TGB2ATGF | TGB2KZTGR |
| **Amino-acid substitution Rluc mutants** | **M1A** | 35S-Rluc | M1AF | 35S-up1R |
| | **M14A** | 35S-Rluc | M14AF | 35S-up1R |
| | **M14F** | 35S-Rluc | M14FF | 35S-up1R |
| | **M14L** | 35S-Rluc | M14LF | 35S-up1R |
| | **M14P** | 35S-Rluc | M14PF | 35S-up1R |
| | **M14S** | 35S-Rluc | M14SF | 35S-up1R |
| | **M14T** | 35S-Rluc | M14TF | 35S-up1R |
| | **M14V** | 35S-Rluc | M14VF | 35S-up1R |
| | **M1A_M14A** | M1A | M14AF | M14R |
| | **M1A_M14F** | M1A | M14FF | M14R |
| | **M1A_M14L** | M1A | M14LF | M14R |
| | **M1A_M14T** | M1A | M14TF | M14R |
| | **M1A_M14V** | M1A | M14VF | M14R |
| **Leader sequence variants for agroinoculation** | **PlAMV-sgRNA1-5U1nt** | pPlAMV | 35S-1ntF | GRR |
| | **PlAMV-sgRNA1-5U3nt** | pPlAMV | 35S-3ntF | GRR |
| | **PlAMV-sgRNA1-5U5nt** | pPlAMV | 35S-5ntF | GRR |
| | **PlAMV-sgRNA1-5U7nt** | pPlAMV | 35S-7ntF | GRR |
| | **PlAMV-sgRNA1-5U10nt** | pPlAMV | 35S-10ntF | GRR |
| | **PVX-sgRNA1-1nt** | pPVX | 35S-PVX1ntF | GRR |
| | **PVX-sgRNA1-4nt** | pPVX | 35S-PVX4ntF | GRR |
| | **PVX-sgRNA1-10nt** | pPVX | 35S-PVX10ntF | GRR |
| | **PVX-GFP-5U4nt** | pPVX-GFP | 35SF | PVX4ntR |
| | **PVX-GFP-5U10nt** | pPVX-GFP | 36SF | PVX10ntR |

Supplementary Table 5

```
#! /usr/local/bin/perl

use strict;
use warnings;
use Getopt::Std;

#  perl parse-genbank1.pl -i folder name -o result file name
#  Read the genbank file in the specified folder
#  Output the position of the start of the mRNA and the position of the first ATG to appear.
#  The result file will be created in the specified folder.

my $mode = 0;
my $name = "";
my $sep = "¥t";
my %opts;
my @list;

getopts('i:o:', ¥%opts);
my $in = $opts{'i'} or die "use: $0 -i infile¥n";
my $out = $opts{'o'} or die "use: $0 -o outfile¥n";

opendir(DIR,"$in");
		my @data = readdir(DIR);

		chdir $in;
		open(OUT,">$out");

print OUT "LOCUS",$sep,"START",$sep,"STOP",$sep,"ATG_pos",$sep,
		"sequence_before_ATG","¥n";

foreach my $file (@data) {
		if ($file eq "." or $file eq "..") {next}
		open(IN,"<$file");
		$mode = 0;

while (<IN>) {

		if ($mode == 0 and /^LOCUS¥s+(.+?)¥s+/) {
				print OUT $1;
				$mode = 1;
		}

		if ($mode == 1 and /^¥s+CDS¥s+(¥d+)¥.¥.(¥d+)/) {
				$mode = 2;
				print OUT $sep,$1,$sep,$2;
		}

		if ($mode == 3 and /^¥/¥//) {
				$mode = 4;
				@list = split(/(atg)/,$name);
				print OUT $sep,length($list[0])+1,$sep,$list[0];
		}

		if ($mode == 3) {
				chomp;
				s/¥s//g;
				s/¥d//g;
				$name .= $_;
		}

		if ($mode == 2 and /^ORIGIN/) {
				$mode = 3;
				$name = "";
		}

}
		print OUT "¥n";
		close IN;
}

		close OUT;
closedir(DIR);
```
